# Mapping circuit dynamics during function and dysfunction

**DOI:** 10.1101/2021.07.06.451370

**Authors:** Srinivas Gorur-Shandilya, Elizabeth M. Cronin, Anna C. Schneider, Sara Ann Haddad, Philipp Rosenbaum, Dirk Bucher, Farzan Nadim, Eve Marder

**Affiliations:** Volen Center and Biology Department, Brandeis University, Waltham MA 02454 USA; Federated Department of Biological Sciences, New Jersey Institute of Technology and Rutgers University, 323 Martin Luther King Blvd, Newark, NJ 07102

## Abstract

Neural circuits can generate many spike patterns, but only some are functional. The study of how circuits generate and maintain functional dynamics is hindered by a poverty of description of circuit dynamics across functional and dysfunctional states. For example, although the regular oscillation of a central pattern generator is well characterized by its frequency and the phase relationships between its neurons, these metrics are ineffective descriptors of the irregular and aperiodic dynamics that circuits can generate under perturbation or in disease states. By recording the circuit dynamics of the well-studied pyloric circuit in *C. borealis*, we used statistical features of spike times from neurons in the circuit to visualize the spike patterns generated by this circuit under a variety of conditions. This unsupervised approach captures both the variability of functional rhythms and the diversity of atypical dynamics in a single map. Clusters in the map identify qualitatively different spike patterns hinting at different dynamical states in the circuit. State probability and the statistics of the transitions between states varied with environmental perturbations, removal of descending neuromodulation, and the addition of exogenous neuromodulators. This analysis reveals strong mechanistically interpretable links between complex changes in the collective behavior of a neural circuit and specific experimental manipulations, and can constrain hypotheses of how circuits generate functional dynamics despite variability in circuit architecture and environmental perturbations.

## Introduction

Neural circuits can generate a wide variety of spiking dynamics, but must constrain their dynamics to function appropriately. Cortical circuits maintain irregular spiking patterns through a balance of excitatory and inhibitory inputs (van Vreeswijk and Sompolinsky, 1996; Mariño et al., 2005; Brunel and Wang, 2003) and the loss of canonical dynamics is associated with neural diseases like channelopathies and epilepsy (Marbán, 2002; Staley, 2015). Preserving functional dynamics can be a challenge for neural circuits for the following reasons. The same spike pattern can be generated by diverse circuits with many different topologies and broadly distributed synaptic and cellular parameters (Prinz et al., 2004; Golowasch et al., 2002; Alonso and Marder, 2019). Furthermore, neural circuits are constantly being reconfigured, with ion channel protein turnover, and homeostatic feedback mechanisms modifying conductance and synapse strengths continuously (Turrigiano et al., 1994, 1995; O’Leary et al., 2014; Franci et al., 2020). The problem of maintaining functional activity patterns is aggravated by the fact that functional circuit dynamics tend to lie within a low-dimensional subspace within the high-dimensional state space: of the numerous possible solutions, only a few are functional and are found in animals (Cunningham and Yu, 2014; Pang et al., 2016). How do neural circuits preserve functional dynamics despite these obstacles?

Answering this question requires, as a prerequisite, a quantitative description of the dynamics of neural circuits during function and dysfunction. When rhythms are regular, this is relatively simple, but when rhythms become irregular, classifying them becomes hard (Haddad and Marder, 2018; Tang et al., 2012; Haley et al., 2018). In this paper we study the dynamics of a well-studied central pattern generator, the pyloric circuit in the stomatogastric ganglion in *C. borealis* (Marder and Bucher, 2007). The pyloric circuit is small, in crabs consisting of 13 neurons coupled by inhibitory and electrical synapses. Its topology and cellular dynamics are well understood, and the circuit generates a clearly defined “functional” collective behavior where bursts of spikes from three different cell types alternate rhythmically to generate a triphasic motor pattern. The stereotypy and periodicity of the motor pattern suggests that the dynamics of the pyloric circuit are fundamentally low dimensional. This has allowed for the effective parameterization of the rhythm by a small number of *ad-hoc* descriptors such as the burst period, duty cycles, and phase of each neuron (Hartline and Maynard, 1975; Eisen and Marder, 1984; Miller and Selverston, 1982).

In response to prolonged perturbations, pyloric circuit dynamics are not always periodic, and descriptors that work well to characterize the canonical rhythm are inadequate to describe these atypical dynamical states. Efforts to study circuit dynamics under these regimes, and to characterize how the circuit responds to, and recovers from perturbations, have been frustrated by the inability to quantitatively describe irregular and non-stationary dynamics (Haddad and Marder, 2018; Tang et al., 2012; Haley et al., 2018).

In this paper we set out to address the problem of quantitatively describing neural circuit dynamics under a variety of conditions. We reasoned that circuit dynamics lie on some lower dimensional set within the full high dimensional space of possible dynamics, even when circuits exhibit atypical and non-functional behavior, because even circuits generating dysfunctional dynamics are still constrained by cellular parameters and network topology. We therefore set out to find and visualize this subset of spike patterns using an unsupervised machine learning approach. This unsupervised method allows us to visualize the totality of a large and complex data set of spike patterns, while being explicit about the assumptions and biases in the analysis. Using this method, we found non-trivial spiking patterns in the distribution of the data that hinted at diverse, stereotyped responses to perturbations. By classifying these patterns, and measuring transitions between these patterns, we were able to characterize the diversity of circuit dynamics under baseline and perturbed conditions, and to identify anecdotally observed atypical states within the full repertoire of spiking patterns (for many hundreds of animals).

## Results

### Perturbations can destabilize the triphasic pyloric rhythm

Studies that measure the pyloric rhythm commonly involve recording from nerves from the stomatogastric ganglion (STG) in ex-vivo preparations. Preparations typically also include the stomatogastric nerve (*stn*) that carries the axons of descending neuromodulatory neurons from the oesophageal and commissural ganglia that project into the STG. Under baseline conditions (11°C, with the *stn* intact, Figure 1a), the periodic triphasic oscillation of the pyloric circuit can be measured by extracellular recordings of the *lpn*, *pdn* and *pyn* nerves (Figure 1a). Bursts of PD spikes on the *pdn* are followed by bursts of LP spikes on *lpn* and bursts of PY spikes on *pyn*. Spikes from LPG neurons are also found on the *pyn* nerve in these recordings, and can be differentiated from PY spikes by their shape and their timing (LPG spikes during PD bursts). Under these control conditions, where the rhythm is robust and spikes from these neurons are easily identifiable both by their location on the nerve and their phase in the cycle, the dual problems of identifying spikes from raw extracellular recordings and meaningfully describing circuit dynamics is easily resolvable.

**Figure 1.**
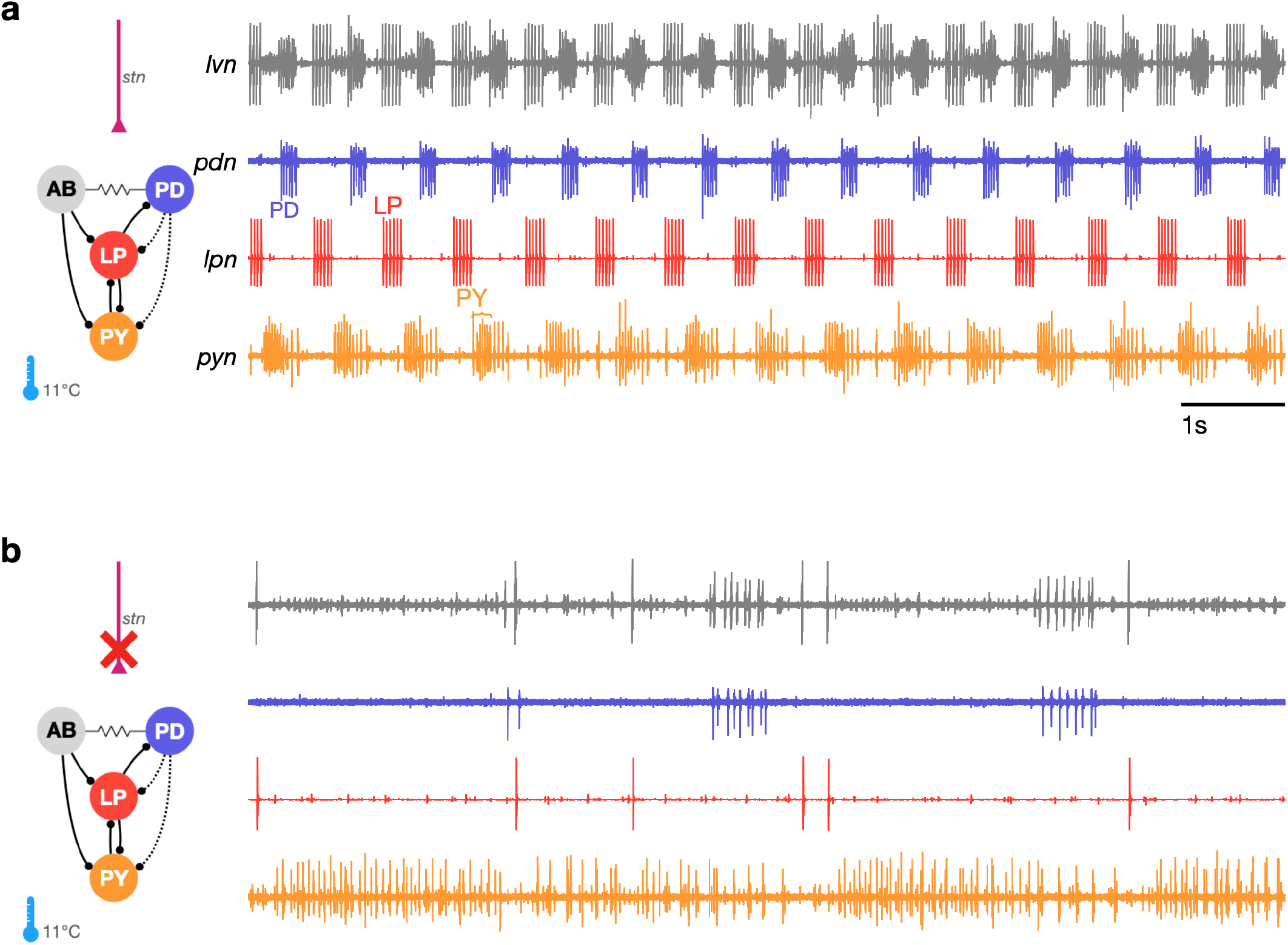
The triphasic pyloric rhythm can become irregular and hard to characterize under perturbation. (a) Simplified schematic of part of the pyloric circuit (left). Filled circles indicate inhibitory synapses. Solid lines are glutamatergic synapses and dotted lines are cholinergic synapses. Resistor symbol indicates electrical coupling. The pyloric circuit is subject to descending neuromodulatory control from the stomatogastric nerve (*stn*). (Right) simultaneous extracellular recordings from the *lvn*, *lpn*, *pdn* and *pyn* motor nerves. Action potentials from LP, PD and PY are visible on *lpn*, *pdn* and *pyn*. Under these baseline conditions, PD, LP and PY neurons burst in a triphasic pattern. The AB neuron is an endogenous burster and is electrically coupled to PD neurons. (b) When the *stn* is cut, neuromodulatory input is removed and the circuit is “decentralized”. In this case, the pyloric rhythm can become irregular and hard to characterize. In addition, spikes from multiple PY neurons can become harder to reliably identify on *pyn*.

In studies that characterize the changes in circuit dynamics to prolonged perturbations, spike identification and circuit dynamics characterization is less straightforward. For example, when descending neuromodulatory projections from the *stn* are cut (i.e., when the STG is decentralized, Figure 1b), the collective dynamics of the pyloric circuit can become less regular. This loss of regularity is concomitant with spikes being harder to reliably identify in extracellular recordings. While PD and LP neuron spikes can still be typically easily identified on the *pdn* and *lpn* nerves (Figure 1b), identifying PY on the *pyn* in the absence of a regular rhythm can be challenging. This problem is aggravated by the fact that spikes from the LPG neuron are frequently found on *pyn*, and because there are several copies of the PY neuron, whose spikes can range from perfect coincidence to slight offsets that can unpredictably change the amplitude and shape of PY spikes due to partial summation. For these reasons, some previous work studying the response of pyloric circuits to perturbations have consistently recorded from the *lpn* and *pdn* nerves, but not from the *pyn* (Hamood et al., 2015; Haley et al., 2018; Haddad and Marder, 2018; Rosenbaum and Marder, 2018). Therefore, in order to include the largest number of experiments in our meta-analysis, we chose to characterize the dynamics of the LP and PD neurons.

### Nonlinear dimensionality reduction allows for the visualization of diverse pyloric circuit dynamics

The regular pyloric rhythm involves out-of-phase bursts of spikes between LP and PD, and is observed under baseline conditions (Figure 2a 1-3). Perturbations such as the removal of descending neuromodulatory inputs, changes in temperature, or changes in pH can qualitatively alter the rhythm, leading to a large variety of hard-to-characterize spiking patterns (Figure 2a 4-6). Because these irregular states may lose the strong periodicity found in the canonical motor pattern, burst metrics such as burst period or phase offsets between bursts that work well to characterize the regular rhythm perform poorly. Efforts to characterize and quantify these atypical spike patterns must overcome the slow timescales in observed dynamics, the large quantity of data, and irregularity and variability in observed spike trains. Previous work used *ad-hoc* categorization systems to assign observations of spike trains into one of a few groups (Haddad and Marder, 2018; Haley et al., 2018), but these categorization methods scaled poorly and relied on subjective annotations.

**Figure 2.**
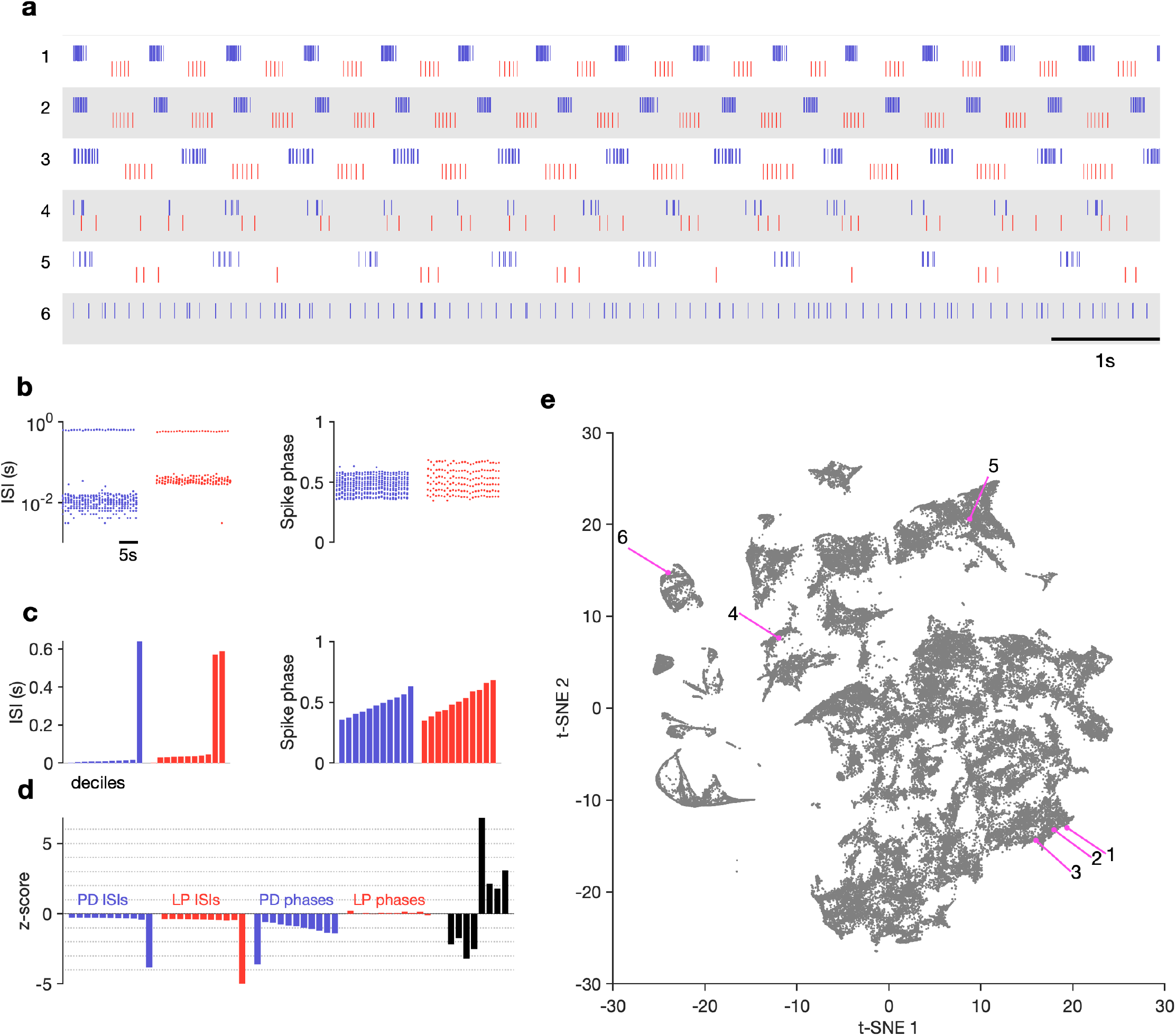
Visualization of diverse neural circuit dynamics. (a) Examples of canonical (1-3) and atypical (4-6) spike patterns of PD (blue) and LP (red). Rasters show 10s of data. (b-d) Schematic of data analysis pipeline. (b) Spike rasters in (a-2) can be equivalently represented by inter-spike intervals and phases. (c) Summary statistics of ISI and phase sets in (d), showing tenth-percentiles. (d) *z*-scored data assembled into a single vector, together with some additional measures (Methods and Materials). (e) Embedding of data matrix containing all vectors such as the one shown in (d) using t-SNE. Each dot in this image corresponds to a single 20-second spike train from both LP and PD. Example spike patterns shown in (a) are highlighted in the map. *n* = 94844 points from *N* = 426 animals. In (a-d), features derived from with LP spike times are shown in red, and features derived from PD spike times are shown in blue. **Figure 2-Figure supplement 1.** Burst metrics smoothly vary in map. **Figure 2-Figure supplement 2.** Embedding arranges data so that neighbors tend to be similar. **Figure 2-Figure supplement 3.** Effect of varying perplexity in t-SNE embedding.

We sought instead to visualize the totality of pyloric circuit dynamics under all conditions using an unsupervised method that did not rely on *a-priori* identification of canonical dynamical patterns. Such a data visualization method, while descriptive, would generate a quantitative vocabulary to catalogue the diversity of spike patterns observed both when these patterns were regular and also when they were irregular and aperiodic, thus allowing for the quantitative characterization of data previously inaccessible to traditional methods (Börner et al., 2003; Nguyen and Holmes, 2019).

The visualization was generated as follows: time-binned (20s) spike trains were converted into their equivalent inter-spike interval (ISI) and phase representations (Figure 2b, Methods and Materials). Because there can be an arbitrary number of spikes in a bin, there are an arbitrary number of ISIs and phases. To convert this into a vector of fixed length, we measured percentiles of ISIs and phases (Figure 2c). Together with other metrics (Methods and Materials), these percentiles were assembled into a fixed-length vector and each dimension was *z*-scored across the entire dataset (Figure 2d). A collection of spike trains from an arbitrary number of neurons has thus been reduced to a matrix where each row consists of *z*-scored percentiles of ISIs and other metrics. This matrix can be visualized using a non-linear dimensionality reduction technique such as *t*-distributed stochastic neighbor embedding (t-SNE) (Van der Maaten and Hinton, 2008), which can generate a two-dimensional representation of the full data set (Figure 2e).

In this representation, each dot corresponds to a single time bin of spike trains from both neurons. We found that spike trains that are visually similar (Figure 2a 1-3) tend to occur close to each other in the embedding (Figure 2e 1-3). Spike patterns that are qualitatively different from each other (Figure 2a4-6) tended to occur far from each other, often in clusters separated by regions of low data density (Figure 2e4-6).

How useful is such a visualization and does it represent the variation in spike patterns in the data in a reasonable manner? We colored each point by classically defined features such as the burst period or the phase (Figure 2–Figure Supplement 1). We found that the embedding arranges data so that differences between clusters and within clusters had interpretable differences in various burst metrics. For example, clusters on the left edge of the map tended not to have defined LP phases, typically due to silent or very sparse LP firing (Figure 2–Figure Supplement 1b). Location of data in the largest cluster was correlated to firing rate in the PD neuron (Figure 2–Figure Supplement 1c). We observed that burst metrics, when they were defined, tended to vary smoothly across the map. To quantify this observation, we built a Delaunay triangulation (Methods and Materials) on the embedded data and measured the triadic differences between PD burst periods and PD duty cycles (Figure 2–Figure Supplement 2). Triadic differences in these metrics were significantly smaller in the map than triadic differences in a projection of the first two principal components or a shuffled map (*p < .*0001, Kolmogorov-Smirnoff test), suggesting that the t-SNE cost function generates a useful embedding where spike features vary smoothly within clusters.

### Visualization of circuit dynamics allows manual labelling and clustering of data

Previous studies have shown that regular oscillatory bursting activity of the pyloric circuit can qualitatively change on perturbation. Circuit dynamics can be highly variable, and has been categorized into various states such as “atypical firing”, “LP-01 spikes” or “atypical” (Haddad and Marder, 2018; Haley et al., 2018). Both the process of constructing these categories and the process of classifying data into these categories are typically done manually, and therefore requires expert knowledge that is not explicitly captured and is impossible to reproduce. Because the embedding distributed data into clusters, we hypothesized that clusters corresponded to stereotyped dynamics that were largely similar, and different clusters represented the qualitatively different circuit dynamics identified by earlier studies.

We therefore inspected circuit dynamics at randomly chosen points in each apparent cluster, and generated labels to describe the dynamics in that region (Figure 3). This process colored the map and segmented it into distinct regions that broadly followed, and were largely determined by, the distribution of the data in the embedding (Figure 3a). Most of the data (57%) were assigned the `regular` label, where both PD and LP neurons burst regularly in alternation with at least two spikes per burst, and all identified `regular` states occurred in a single contiguous region in the map (blue). In the `LP-weak-skipped` state, PD bursts regularly, but LP does not burst every cycle, or only fires a single spike per burst. `irregular-bursting` states showed bursting activity on both neurons, which were interrupted or otherwise irregular. In contrast, the irregular state showed spiking that was more variable, and did not show strong signs of bursting at any point. `LP-silent-PD-bursting` states had regular bursting on PD, with no spikes on LP, while LP-silent states also had no spikes on LP, but activity on PD was more variable, and did not show regular bursting.

**Figure 3.**
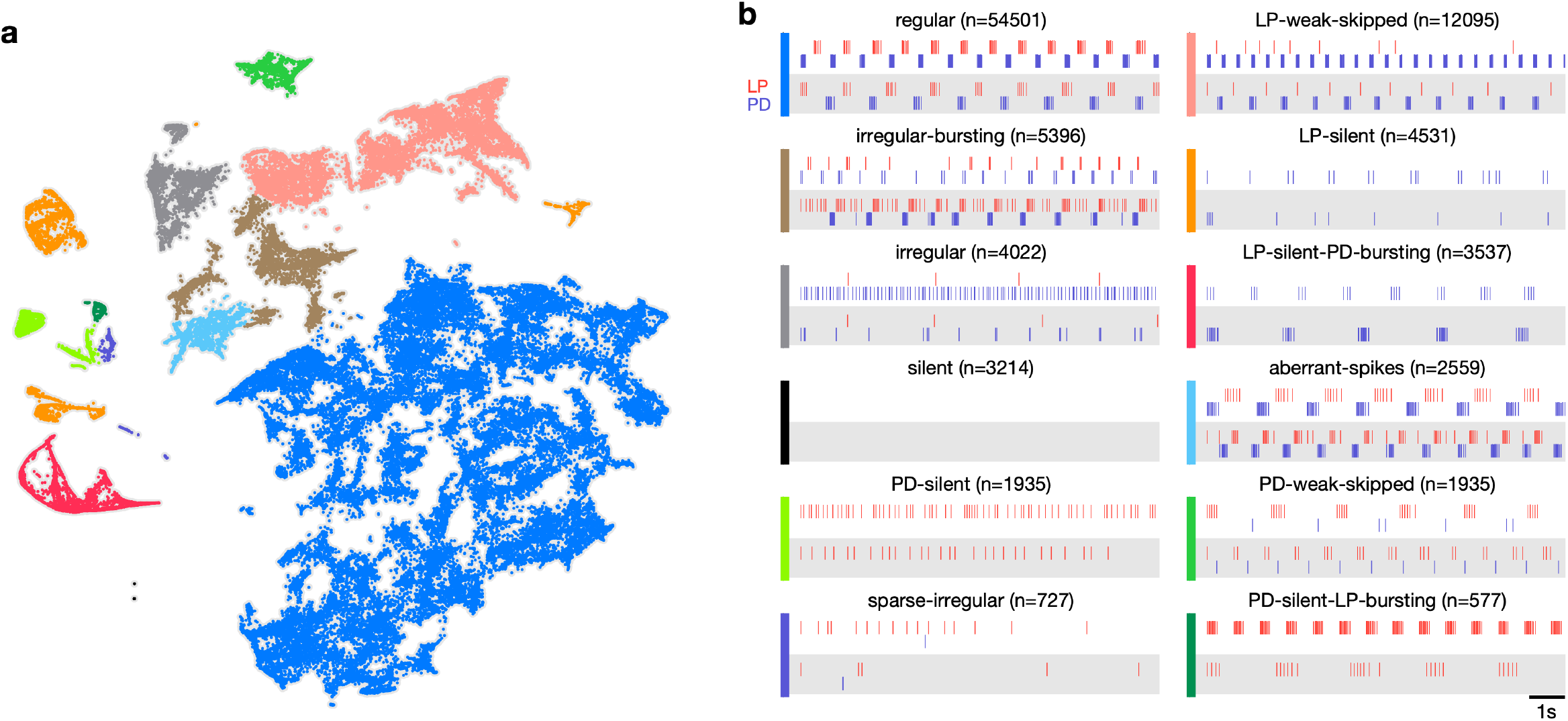
Map allows identification of distinct spiking dynamics. (a) Map of all pyloric dynamics in dataset where each point is colored by manually assigned labels. Each point corresponds to a 20s paired spike train from LP and PD. Each panel in (b) shows two randomly chosen points from that class. The number of points in each class is shown in parentheses above each panel. *n* = 94844 points from *N* = 426 animals. Labels are ordered by likelihood in the data. **Figure 3-Figure supplement 1.** Speed of trajectories through map. **Figure 3-Figure supplement 2.** Embeddings with different initializations.

The time evolution of the pyloric dynamics of every preparation constitutes a trajectory in the map, and every point in the map is therefore associated with an instantaneous speed of motion in the map. We hypothesized that instantaneous speed could vary across the map, with points labelled regular moving more slowly through the map than points with labels corresponding to atypical states such as `irregular`, because `regular` rhythms would vary less over time. Consistent with this, we found that points in the regular cluster tended to have smaller speeds than points in other clusters (Figure 3–Figure Supplement 1a). Speeds in the `regular` state we significantly lower than every other state except `PD-silent-LP-bursting` (*p < .*004, permutation test), suggesting that atypical states were associated with increased variability in circuit dynamics (Figure 3–Figure Supplement 1b).

### Variability in baseline circuit dynamics across a population of wild-caught animals

Work on the pyloric circuit has almost exclusively used a wild-caught crustacean population. This uncontrolled environmental and genetic variability serves as a window into the extant variability of a functional neural circuit in a wild population of animals. In addition, experimental and computational work has shown that similar rhythms can be generated by a wide variety of circuit architectures and cellular parameters (Prinz et al., 2003; Hamood and Marder, 2015; Alonso and Marder, 2019). We therefore set out to study the variability in baseline circuit dynamics in the 346 pyloric circuits recorded from under baseline conditions in this dataset.

The burst period of the pyloric circuit in the lobster can vary 2-3 fold under baseline conditions at 11°C across animals (Bucher et al., 2005). Despite this sizable variation, other burst metrics, such as the phase onset of follower neurons, or the duty cycles of individual neurons, are tightly constrained (Bucher et al., 2005), likely related to the fact that these circuits are under activity-dependent feedback regulation (Turrigiano et al., 1995; O’Leary et al., 2014; Gorur-Shandilya et al., 2020) as they develop and grow. Activity-dependent regulation of diverse pyloric circuits could constrain variability in a single circuit across time to be smaller than variability across the population. To test this hypothesis, we measured a number of burst metrics such as burst period and the phases and duty cycles of the two neurons across these 346 preparations in baseline conditions (Figure 4) when data are labelled regular, because metrics are well-defined in this state. Mean values of each of these metrics were unimodally distributed (Figure 4a) and the coefficient of variation for all metrics was approximately 0.1 (Figure 4b). Using the mean coefficient of variation in each individual as a proxy for the within-animal variability, and the coefficient of variation of the individual means as a proxy for the across-animal variability, we found that every metric measured was more variable across animals than within animals (Figure 4c). Shuffling experimental labels generated null distributions for excess variability across animals, and showed that across animal variability was significantly greater than within animal variability (Figure 4d, *p < .*007, permutation test, Table 1).

**Figure 4.**
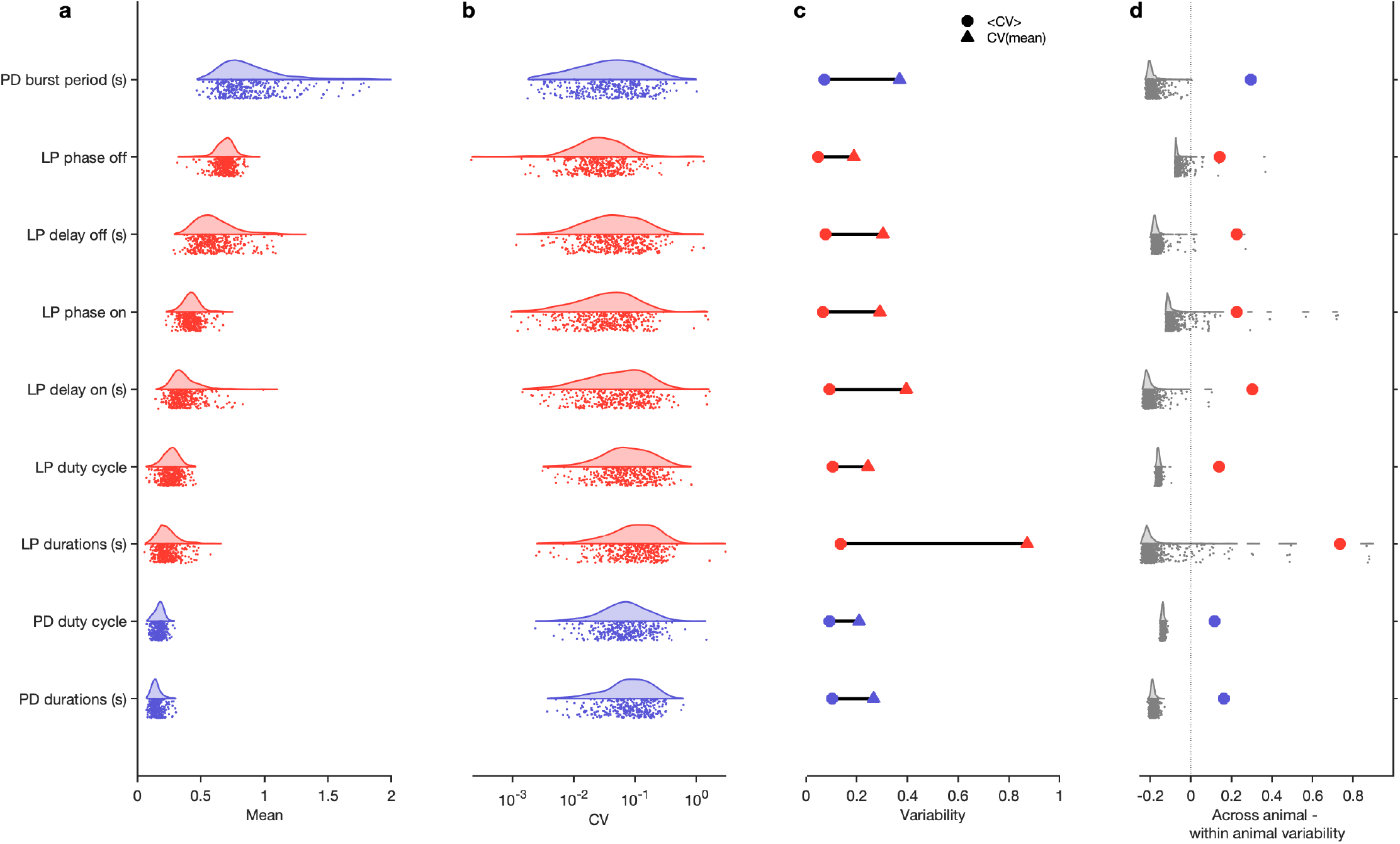
Variability of burst metrics under baseline conditions. (a) Variability of burst metrics in PD and LP neurons across a population of wild caught animals. Metrics are only computed under baseline conditions and in the regular cluster. (b) Distribution of coefficient of variation (CV) of metrics in each animal across all data from that animal. In (a-b), each dot is from a single animal, and distributions show variability across the entire population. (c) Across-animal variability (CV of individual means, Δ) is greater than within-animal variation (mean of CV in each animal, ⃝) for every metric. (d) Difference between across animal variability and within animal variability (colored dots). For each metric, gray dots and distribution show differences between across-animal and within-animal variability for shuffled data. *n* = 18336 points from *N* = 346 animals. **Figure 4-Figure supplement 1.** State distribution under baseline conditions **Figure 4-Figure supplement 2.** Recording condition alters regular state probability **Figure 4-Figure supplement 3.** Effect of sea surface temperature on baseline circuit dynamics

**Table 1.** ANOVA results and power analysis for *Figure 4*. Table 1–source data 1. ANOVA results for burst metrics in baseline conditions. For each metric, each animal is treated as a group and the variability (mean square difference) is compared within and across group. *F* is the ratio of across-animal to within-animal mean square differences. *N.*_99_ is the estimate of the sample size required to reject the null hypothesis with a probability of .99 when the alternative hypothesis is true. *N* = 346 animals.

| Metric | Across animal MS | Within animal MS | F | $N_{.99}$ |
| --- | --- | --- | --- | --- |
| LP delay off (s) | 1.1391 | 0.010956 | 103.97 | 6 |
| LP delay on (s) | 0.61647 | 0.0111 | 55.54 | 6 |
| LP durations (s) | 0.36386 | 0.012366 | 29.424 | 4 |
| LP duty cycle | 0.15986 | 0.0013093 | 122.09 | 10 |
| LP phase off | 0.23406 | 0.0072279 | 32.383 | 11 |
| LP phase on | 0.21655 | 0.0088115 | 24.576 | 9 |
| PD burst period (s) | 3.557 | 0.036872 | 96.469 | 4 |
| PD durations (s) | 0.079397 | 0.00054944 | 144.5 | 6 |
| PD duty cycle | 0.053472 | 0.00041323 | 129.4 | 16 |

It is reasonable to suppose that all baseline data exist in the regular cluster. While most baseline data are confined to the regular cluster (≈80%, Figure 4–Figure Supplement 1a), the remaining data, nominally recorded under baseline conditions, contains atypical circuit dynamics (Figure 4–Figure Supplement 1b-c). What causes these atypical circuit dynamics in this large, unbiased survey of baseline pyloric activity? One possibility could be inadvertent damage to the preparation caused by dissection and preparation of the circuit for recording. Consistent with this, we found that the probability of observing `regular` states was significantly reduced when cells were recorded from intracellularly (Figure 4–Figure Supplement 2), which may be due to increase in leak currents due to impaling cells with sharp electrodes (Cymbalyuk et al., 2002), or due to cell dialysis (Hooper et al., 2015). No significant correlation was observed between sea surface temperatures (a proxy for environmental conditions for these wild-caught animals) and burst metrics (Figure 4–Figure Supplement 4a-c) or the probability of observing a regular state (Figure 4–Figure Supplement 4d). Taken together, these results underscore the importance of verifying that baseline or control data does not include uncontrolled technical variability that could mask biological effects of interest.

### Perturbation modality alters state probability

The pyloric circuit and other circuits in the crab must exhibit robustness to the environmental perturbations that these animals are likely to encounter. Previous studies have characterized the ability of crustacean circuits to be robust to environmental perturbations such as pH (Haley et al., 2018; Ratliff et al., 2021; Qadri et al., 2007), temperature (Tang et al., 2010, 2012; Rinberg et al., 2013; Haddad and Marder, 2018; Kushinsky et al., 2019), oxygen levels (Clemens et al., 2001) and changes in extracellular ionic concentrations (He et al., 2020). Robustness to these perturbations exists up to a limit, likely reflecting the bounds of the natural variation in these quantities that these circuits are evolved to function in. When challenged with extremes of any of these perturbation modalities, the pyloric rhythm breaks down, displaying irregular or aberrant states, and may even cease spiking entirely.

What remains unclear is if extreme perturbations of different modalities share common pathways of destabilizing and disrupting the pyloric rhythm (Ratliff et al., 2021). In principle, these environmental perturbations can disrupt neuron and circuit function in qualitatively different ways: e.g., changes in extracellular potassium concentration can alter the reversal potential of potassium (He et al., 2020) vs. changes in temperature can have varied effects on the timescales and conductances of all ion channels (Tang et al., 2010; Caplan et al., 2014). Because prior work was focussed on studying the limits of robustness, and lacked a detailed quantitative description of irregular behavior, the fine structure of the transition between functional dynamics and silent or “crashed” states remain poorly characterized (Ratliff et al., 2021). We therefore set out to measure how pH, temperature and extracellular potassium perturbations alter circuit state probability.

Where in the map are data under extreme environmental perturbations? Circuit spike patterns under high pH (>9.5), high temperature (>25°C) and high extracellular potassium (2.5 × [*K*^+^]) are distributed across a wide region of the map, spanning both regions in the regular cluster and other non-regular clusters (Figure 5a). Spike patterns observed under high temperature conditions in the regular region were clustered in the lower extremity, in the region containing high firing rates and small burst periods of PD (Figure 2–Figure Supplement 1), consistent with earlier studies showing that elevated temperatures tend to speed up the pyloric rhythm (Tang et al., 2010, 2012). To characterize how environmental perturbations destabilize the pyloric rhythm and increase the variability in observed dynamics, we measured the mean distance travelled in the map by each preparation as a function of the perturbation intensity (Figure 5b). For both pH and temperature perturbations, the mean distance travelled in the map was lowest at baseline conditions (pH 7.8, 11°C) and increased away from these conditions, suggesting that changes in either of these environmental parameters increased the variability in observed pyloric dynamics (*ρ* = .95 for pH>7.8, *p* = −.36 for pH<7.8, *ρ* = .81 for T>11°C, *p < .*001, Spearman rank correlation test).

**Figure 5.**
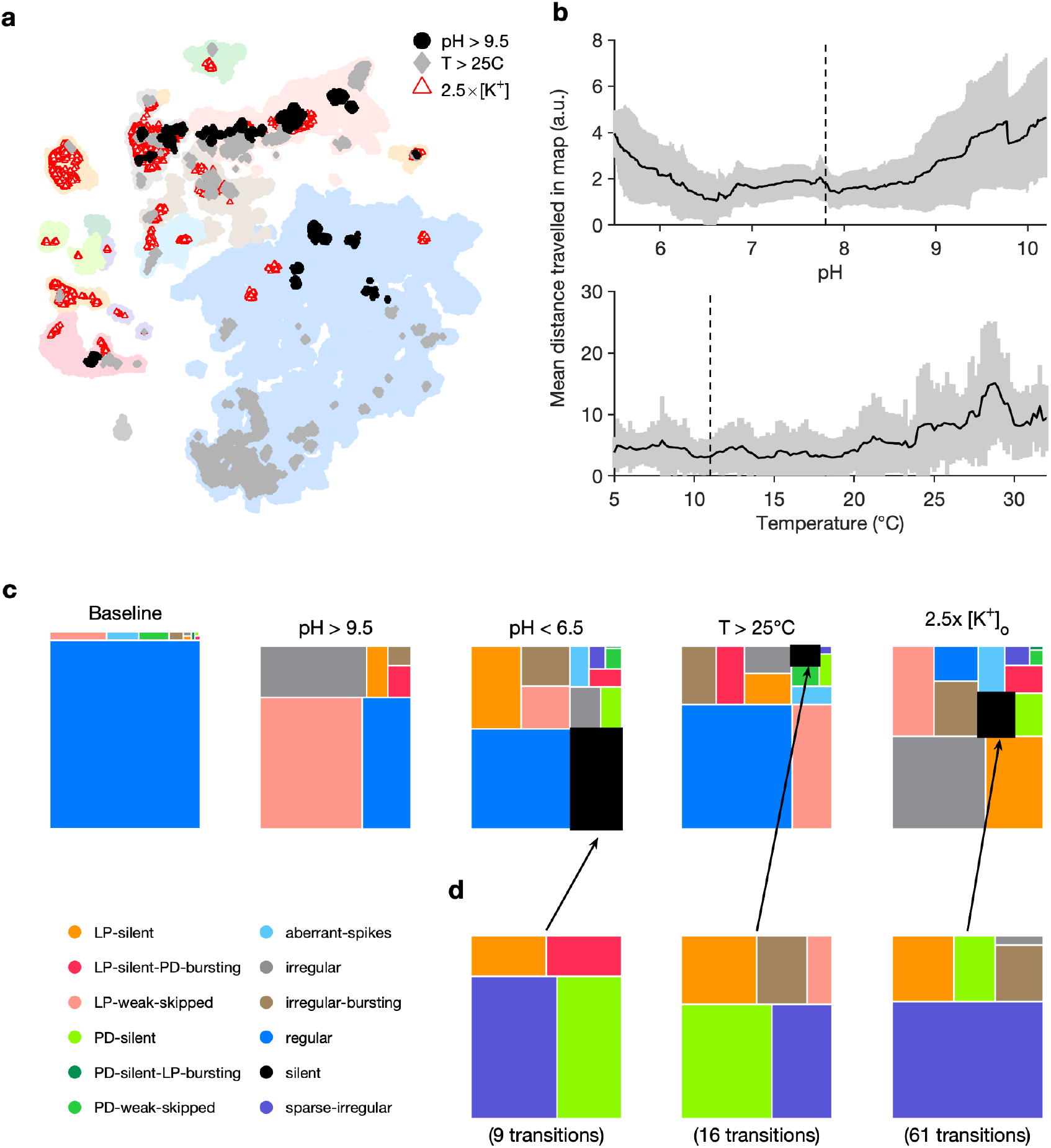
Effect of three different environmental perturbations. (a) Map showing regions that are more likely to contain data recorded under extreme environmental perturbations. (b) Mean distance travelled in map during pH and temperature perturbations. Solid lines indicate mean and shading is the standard deviation across all preparations. Vertical dashed lines indicate baseline conditions. (c) Treemaps showing probability distributions of states under baseline and perturbed conditions. (d) Probability distribution of states preceding silent state under perturbation. pH perturbations: *n* = 4023 from 6 animals. [*K*^+^] perturbations: *n* = 5526 from 20 animals. Temperature perturbations: *n* = 80470 from 414 animals. **Figure 5-Figure supplement 1.** Preparation-by-preparation response to pH perturbations.

Subjecting the pyloric circuit to extremes of pH, temperature and extracellular potassium altered the distribution of observed states (Figure 5c). In all cases, the probability of observing regular was significantly reduced (*p < .*001, paired permutation test), and a variety of non-regular states were observed. We observed that high pH (>9.5) did not silence the preparation, but silent states were observed in low pH (<6.5), consistent with previously published manual annotation of this data (Haley et al., 2018). Silent states were also observed in 2.5 × [*K*^+^], as reported earlier by He et al. (2020). Previous work has shown that the isolated pacemaker kernel (AB and PD neurons) has a stereotyped trajectory from bursting through tonic spiking to silence when subjected to temperature and high [*K*^+^] perturbations (Ratliff et al., 2021). Do pathways to silent states share similarities across perturbation modality in intact circuits? To answer this, we plotted the probability of observing states conditioned on the transition to silence in low pH, high temperature, and 2.5 × [*K*^+^] (Figure 5d). In the ≈ 2000 transitions between states detected, we never observed a transition from regular to silent, suggesting that the timescales of silencing are slow, longer than the width of one data bin (20s). Trajectories to silent states always transition through a few intermediate states such as `sparse-irregular, LP-silent` or `PD-silent` (Figure 5d).

### Transitions between states during environmental perturbations

Changes in temperature, pH and [*K*^+^] have different effects on the cells in the pyloric circuit and therefore can destabilize the rhythm in different ways. Increasing the extracellular [*K*^+^] changes the reversal potential of *K*^+^ ions, altering the currents flowing through potassium channels, and typically depolarizes the neuron (He et al., 2020). Ion channels can be differentially sensitive to changes in temperature or pH, and changes in these variables can have complex effects of ionic currents in neurons (Tang et al., 2010, 2012; Haley et al., 2018). We therefore asked if different environmental perturbations changed the way in which regular rhythms destabilized.

Our analysis mapped a time series of spiketimes from PD and LP neurons to a categorical time series of labels such as regular. We therefore could measure the transitions between states during different environmental perturbations (Methods and Materials). We found that transition matrices between states shared commonalities across environmental perturbations (Figure 6a), such as likely transitions between `regular` and `LP-weak-skipped` states. `PD-silent-LP-bursting` states tended to be followed by `PD-silent` states, in which the LP neuron is spiking, but not bursting regularly. The LP neuron becomes less regular in both transitions, contributing to the loss of regular rhythms. We never observed a transition from regular rhythms `LP-silent` or `PD-silent` states, suggesting slow (>20s) timescales of rhythm collapse. In high pH, every transition away from the regular state was to the `LP-weak-skipped` state, hinting at increased sensitivity of the LP neuron to high pH. High pH perturbations also never silenced the circuit, as previously reported (Haley et al., 2018), and showed fewer and less varied transitions than other perturbations. Are some transitions over- or under-represented in the transition matrix? To determine this, we constructed a null model where transitions occurred with probabilities that scaled with the marginal probability of final states (Methods and Materials). Transitions that occurred significantly more often than predicted by the null model are shown with black borders and those that occurred significantly less often than predicted are shown with filled circles (Figure 6a). Transitions that never occurred, but occurred at significantly non-zero rates in the null model are indicated with diamonds.

**Figure 6.**
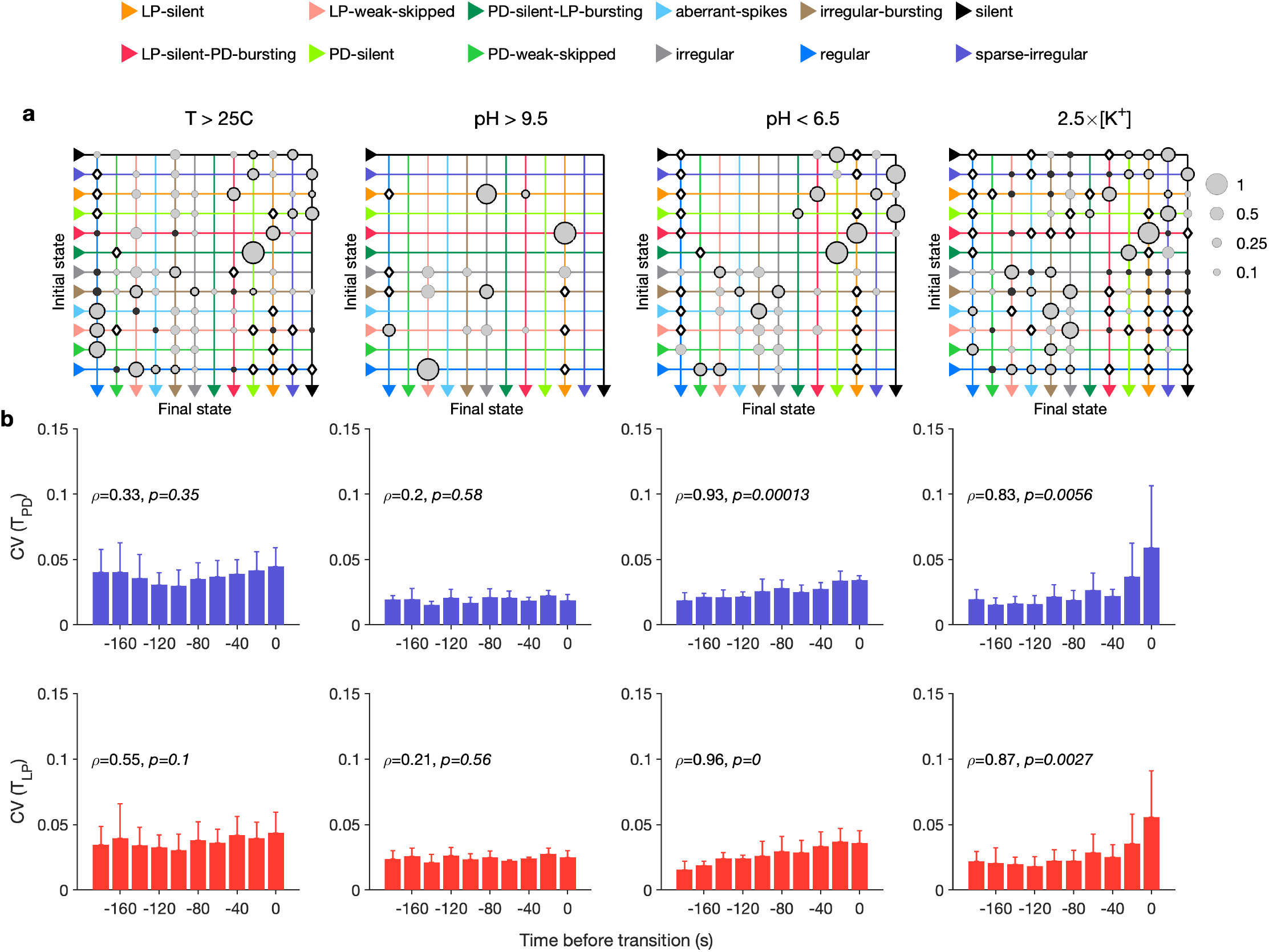
Effect of environmental perturbations on transitions between states. (a) Transition matrix between states during environmental perturbations. Each matrix shows the conditional probability of observing the final state in the next time step given an observation of the initial state. Probabilities in each row sum to 1. Size of disc scales with probability. Discs with dark borders are transitions that are significantly more likely than the null model (Methods and Materials). Dark solid discs are transitions with non-zero probability that are significantly less likely than in the null model. O are transitions that are never observed, and are significantly less likely than in the null model. States are ordered from regular to silent. (b) Coefficient of variation (CV) of burst period of PD (purple) and LP (red) vs. time before transition away from the regular state. *ρ, p* are from Spearman test to check if variability increases significantly before transition. Temperature perturbations: *n* = 1035 transitions in 61 animals. pH perturbations: *n* = 90 transitions in 6 animals. [*K*^+^] perturbations: *n* = 271 transitions in 20 animals.

Earlier work has shown that transitions from regular bursting are preceded by an increase in variability in the voltage dynamics of bursting in PD neurons pharmacologically isolated from most of the pyloric circuit (Ratliff et al., 2021). Can we detect similar signatures of destabilization before transitions from `regular` states in the intact circuit? We measures the coefficient of variation (CV) of the burst periods of PD and LP neurons in `regular` states just before transitions away from regular Figure 6b). Because we restricted our measurement of variability to `regular` states, we could disambiguate true cycle-to-cycle jitter in the timing of bursts from the apparent variability in cycle period due to alternations between bursting and non-bursting dynamics. We found that transitions away from regular were correlated with a steady and almost monotonic increase in variability in PD and LP burst periods for low pH and high [*K*^+^] perturbations, but not for high pH and high temperature perturbations (Spearman rank correlation test). This suggests mechanistically different underpinnings to the pathways of destabilization between these sets of perturbations, and is consistent with previous work showing that robustness to perturbations in pH only moderately affects temperature robustness in the same neuron (Ratliff et al., 2021).

### Decentralization elicits variable circuit dynamics

The pyloric circuit is modulated by a large and chemically diverse family of neuromodulators that it receives via the stomatogastric (*stn*) nerve (Marder, 2012). Decentralization, or the removal of this neuromodulatory input via transection and/or chemical block of the *stn*, has been shown to affect the pyloric rhythm in a number of ways (Russell, 1976). Decentralization can stop the rhythm temporarily, which can recover after a few days (Golowasch et al., 1999; Thoby-Brisson and Simmers, 1998). Decentralization slows down the pyloric rhythm (Eisen and Marder, 1982; Rosenbaum and Marder, 2018), and makes the rhythm more variable (Hamood and Marder, 2015; Hamood et al., 2015). Decentralization can evoke variable circuit dynamics, sometimes with slow timescales (Figure 7–Figure Supplement 1), and can lead to changes in ion channel expression (Mizrahi et al., 2001).

**Figure 7.**
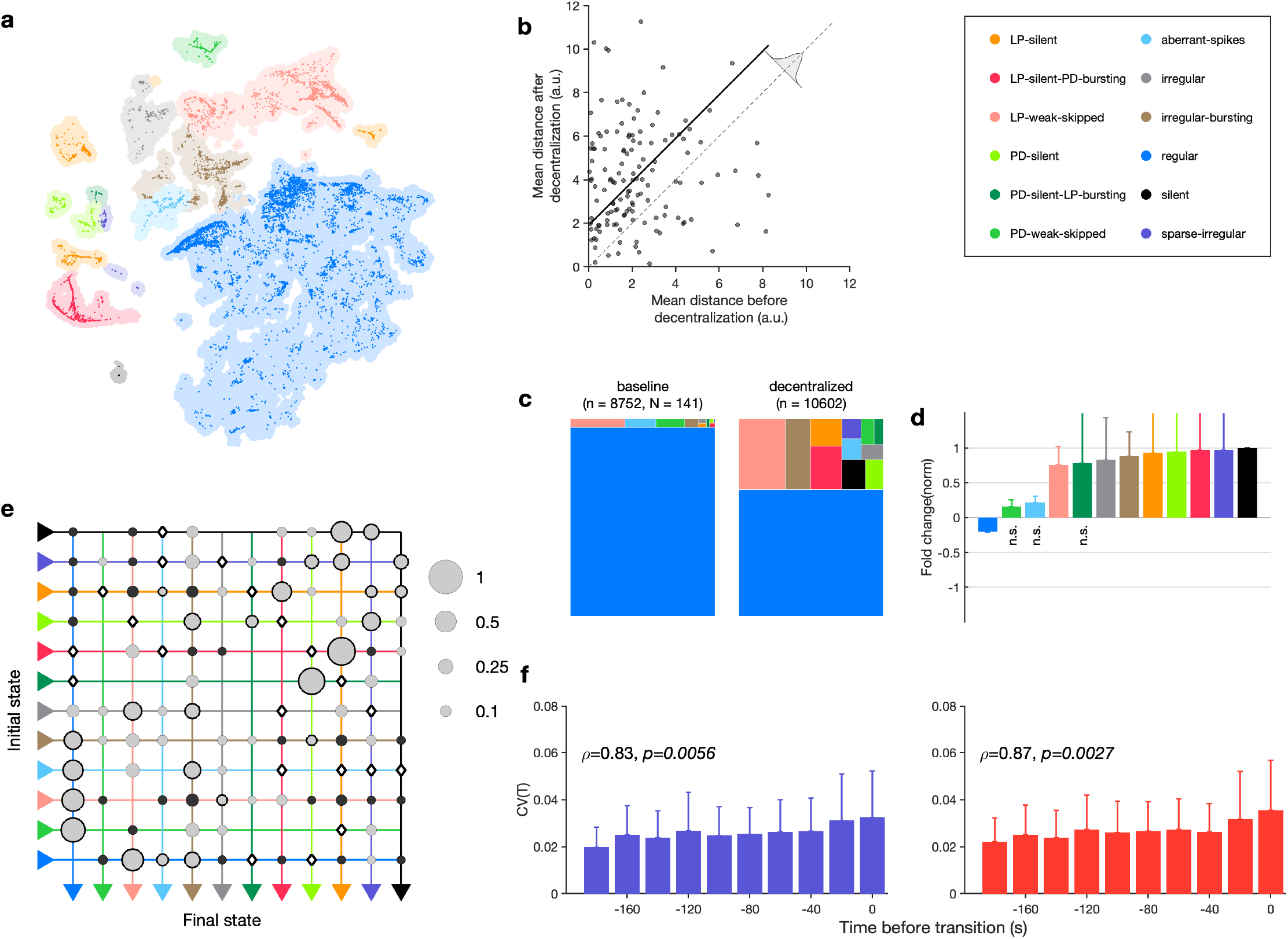
Effect of decentralization. (a) Map occupancy conditional on decentralization. Shading shows all data, bright colored dots indicate data when preparations are decentralized. (b) Distance travelled in map before and after decentralization. Each dot is a single preparation. Gray shading indicates null distribution and solid line is the mean difference upon decentralization. (c) State probabilities before and after decentralization. (d) Fold change in state probabilities on decentralization. States marked n.s. are not significantly more or less likely after decentralization. All other states are (paired permutation test, *p <* 0.00016). (a-b) *n* = 10602 points from *N* = 141 animals. (e) Transition matrix during decentralization. Probabilities in each row sum to 1. Size of disc scales with probability. Discs with dark borders are transitions that are significantly more likely than the null model (Methods and Materials). Dark solid discs are transitions with non-zero probability that are significantly less likely than in the null model. O are transitions that are never observed, and are significantly less likely than in the null model. States are ordered from regular to silent. *n* = 1933 transitions. (f) Coefficient of variation of PD (purple) and LP (red) burst periods before transition away from `regular` states. *p, p* from Spearman test. *n* = 1332 points from *N* = 79 animals. **Figure 7-Figure supplement 1.** Decentralization evokes variable dynamics **Figure 7-Figure supplement 2.** Effects of decentralization on state probabilities **Figure 7-Figure supplement 3.** Time course of effects of decentralization **Figure 7-Figure supplement 4.** Effects of decentralization do not correlate with seasonal effects

The variability in circuit dynamics elicited by decentralization, and the animal-to-animal variability in response to decentralization has made a quantitative analysis of the effects of decentralization difficult. We therefore set about to characterize the variable and invariant features of the changes in circuit spiking dynamics on removal of descending neuromodulation across a large (*N* = 141) population.

We first asked where in the map decentralized data were (Figure 7a). A large fraction (≈ 30%) of the data was found outside the regular cluster, suggesting the existence of atypical circuit dynamics on decentralization. To determine if decentralization dispersed data in the map, and made circuit dynamics more variable across time, we measured the mean distance travelled by each preparation before and after decentralization (Figure 7b, Methods and Materials). Decentralization significantly increased the distance covered by each preparation across the map (*p < .*0001, paired permutation test), suggesting that circuits displayed more variable dynamics on decentralization. Decentralization also changed probabilities of observing many states. The regular state was significantly less likely on decentralization, and several atypical states were significantly more likely (Figure 7c,d, Table 2, Figure 7–Figure Supplement 2).

**Table 2.** State counts before and after decentralization for data shown in Figure 7 Table 2–source data 1. State counts before and after decentralization. *p*-value of change in probability of observing change estimated from paired permutation tests.

| State | $n_{control}$ | $n_{dec.}$ | $p$ | $\Delta P(state)$ |
| --- | --- | --- | --- | --- |
| regular | 7967 | 5791 | <0.001 | -0.308 77 |
| LP-silent | 22 | 724 | <0.001 | 0.030 65 |
| LP-silent-PD-bursting | 14 | 577 | <0.001 | 0.045 926 |
| PD-silent | 11 | 140 | $4 \times 10^{-5}$ | 0.018 51 |
| PD-silent-LP-bursting | 20 | 18 | 0.469 59 | 0.000 188 91 |
| aberrant-spikes | 111 | 168 | 0.300 37 | 0.003 285 3 |
| LP-weak-skipped | 317 | 1628 | <0.001 | 0.099 875 |
| PD-weak-skipped | 142 | 118 | 0.292 19 | 0.003 453 8 |
| sparse-irregular | 4 | 154 | <0.001 | 0.013 263 |
| irregular | 13 | 116 | 0.000 23 | 0.010 877 |
| silent | 0 | 321 | <0.001 | 0.024 825 |
| irregular-bursting | 72 | 753 | <0.001 | 0.057 913 |

How do preparations switch between different states when decentralized? The transition matrix during decentralization revealed many transitions between diverse states (Figure 7e), with the most likely transitions being significantly over-represented compared to the null model (*p < .*05, Methods and Materials). Transitions away from regular included significantly more likely transitions into states where one of the neurons was irregular such as LP-weak-skipped and PD-weak-skipped. Similar to rhythm destabilization in high [*K*^+^] or low pH, transitions away from regular were associated with a near-monotonic increase in the variability of PD and LP burst periods before the transitions (Figure 7f, *ρ* ≈ .8*, p < .*006, Spearman rank correlation test).

The time series of identified states on a preparation-by-preparation basis showed striking variability in the responses to decentralization (Figure 7–Figure Supplement 3a), with the probability of observing `regular` states decreasing immediately after decentral8ization (Figure 7–Figure Supplement 3b). What causes the observed animal-to-animal variability in circuit dynamics on decentralization? One possibility is that seasonal changes in environmental conditions alter the sensitivity of the pyloric circuit to neuromodulation. We tested this hypothesis by measuring the correlation between measures such as the probability of observing the regular state, the change in burst period, and the change in firing rate on decentralization and the sea surface temperature at the approximate location of these wild caught animals (Figure 7–Figure Supplement 4). None of these measures was significantly correlated with sea surface temperature (*p > .*07, Spearman rank correlation test).

### Stereotyped effects of decentralization on burst metrics

Despite the animal-to-animal variation in responses to decentralization, are there stereotyped responses to decentralization? Previous work has shown that decentralization typically slows down the pyloric rhythm (Eisen and Marder, 1982; Rosenbaum and Marder, 2018), but a finer-grained analysis of rhythm metrics were confounded by the irregular dynamics that can arise when preparations are decentralized. For example, alteration between regular and atypical states could bias estimates of burst metrics that are not defined in atypical states. Because our analysis allows us to identify the subset of data where pyloric circuit dynamics are regular enough that burst metrics are well-defined, we measured the changes in a number of burst metrics like the burst period, duty cycle and phases on decentralization (Figure 8a). Every metric measured was significantly changed except the phase at which LP bursts start (*p <* 0.007, paired permutation test). Consistent with earlier studies, we found that the coefficient of variation in every metric increased following decentralization (Figure 8b).

**Figure 8.**
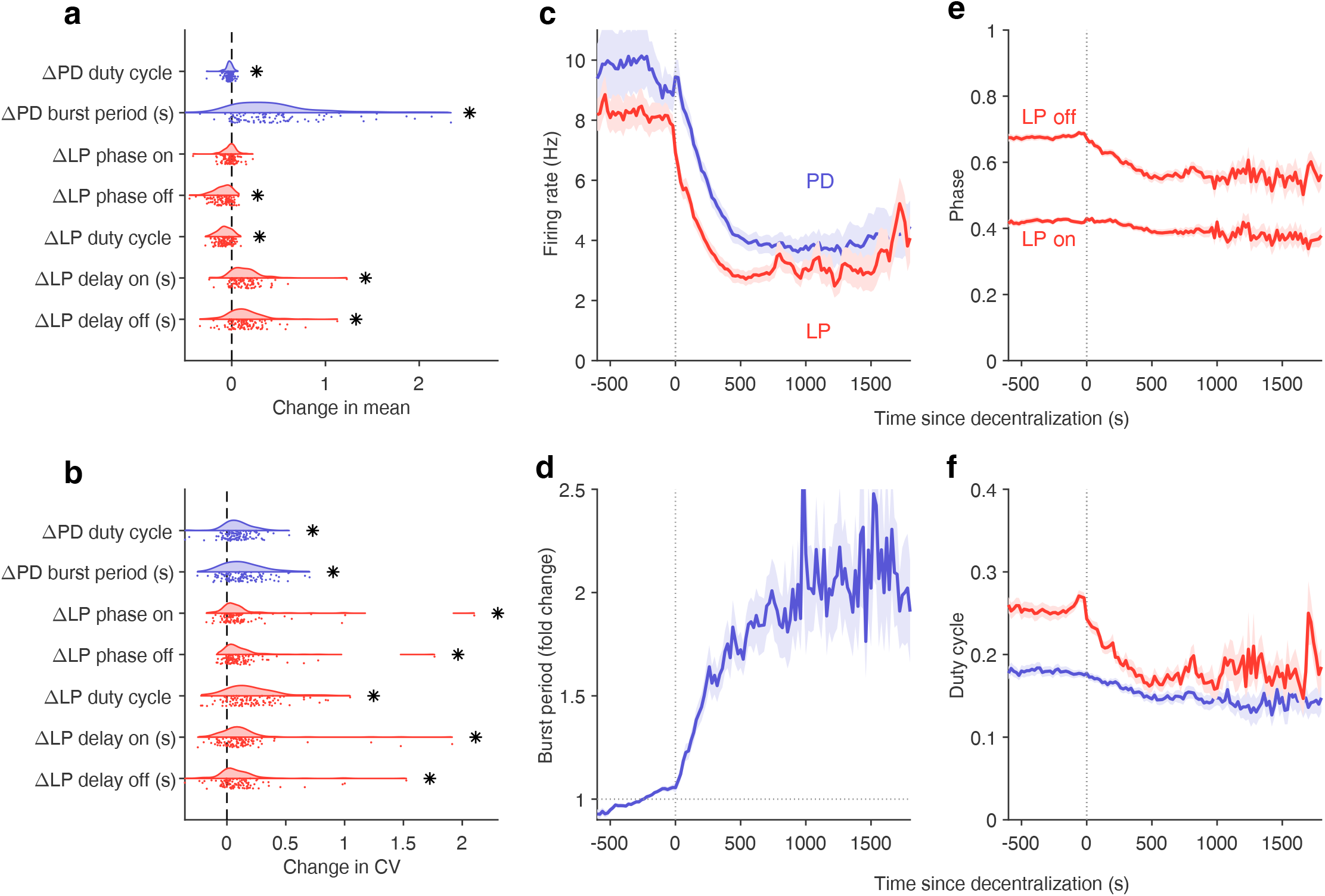
Effects of decentralization on burst metrics. (a) Change in mean burst metrics on decentralization. (b) Change in coefficient of variation of burst metrics on decentralization. In (a) and (b), each dot is a single preparation; * indicate distributions whose mean is significantly different from zero (*p < .*007, paired permutation test). Firing rates (c), burst period (d), LP phases (e) and duty cycles (f) vs. time since decentralization. In panels (c-f), thick lines indicate population means, and shading indicates the standard error of the mean. *n* = 13898 points from *N* = 141 preparations. **Figure 8-Figure supplement 1.** Effects of decentralization on regular rhythms

What are the dynamics of changes in burst metrics on decentralization? Firing rates of both LP and PD neurons decreased immediately on decentralization, roughly halving their pre-decentralized values (Figure 8c). This occurred together with a doubling of PD burst periods (Figure 8d), suggesting that the entire rhythm is slowing down. Intriguingly, decentralization led to significant advance in the phase of LP burst ends, but not starts (Figure 8e), leading to a large decrease in the duty cycle of the LP neuron (Figure 8f) that was significantly more than the decrease in PD’s duty cycle (*p <* 10^−8^, paired *t*-test).

The stereotyped slowing of the rhythm on decentralization can also be quantified by looking at the distribution of the data in the regular cluster before and after decentralization (Figure 8–Figure Supplement 1). Data are concentrated in the upper left edge of the regular cluster when decentralized, where burst periods are large and firing rates low (Figure 2–Figure Supplement 1a,c), suggesting that decentralization could elicit a more stereotyped rhythm for circuits that continue to burst regularly, because circuits that do so tend to share a common, slow bursting dynamics. Counter-intuitively, it may appear that regular rhythms in baseline conditions are more variable than regular rhythms after decentralization. To test this hypothesis we measured the dispersion of each preparation in the map (Figure 8–Figure Supplement 1b) before and after decentralization. Dynamics before decentralization were significantly more dispersed in the regular cluster than dynamics after decentralization (Figure 8–Figure Supplement 1c, *p* = .0016, paired *t*-test), because they then tended to be concentrated in the upper-left edge of that cluster. To first approximation, our analysis shows that there are many ways to manifest a regular rhythm under baseline conditions, but regular rhythms on decentralization are typically slow, and stereotyped in comparison.

### Neuromodulators differentially affect state probabilities

The crustacean stomatogastric ganglion is modulated by more than 30 substances (Harris-Warrick and Marder, 1991; Marder, 2012) that tune neuronal properties on an intermediate time scale, between feedback homeostasis and intrinsic cellular properties (Daur et al., 2016). Earlier work has focussed on understanding the effect modulators have on restoring (or destabilizing) the canonical rhythm, in part because the restoration of regular oscillatory dynamics is a dominant feature of neuromodulator action. Other effects that neuromodulators might have on pyloric circuit dynamics are harder to investigate, and are hindered by the difficulty in characterizing circuit dynamics when non-regular. Here we set out to systematically characterize the effects of neuromodulators on dynamical states identified in the full space of circuit behaviors (Figure 3).

We focussed our analysis on the effect of four neuromodulators: Red pigment-concentrating hormone (RPCH), proctolin, oxotremorine, and serotonin. In the experiments analyzed, these neuromodulators were added to decentralized preparations so that endogenous effects of these (and other) neuromodulators were minimized. We therefore first characterized the distribution of states in decentralized preparations where neuromodulators were subsequently added (Figure 9a). RPCH is a neuropeptide that targets a number of cells in the circuit (Nusbaum and Marder, 1988; Swensen and Marder, 2001), and has been shown to increase the number of spikes per burst in PD and LP, (Dickinson et al., 2001; Thirumalai and Marder, 2002) though it has little effect on the pyloric period (Thirumalai et al., 2006). RPCH increased the probability of the regular state, suggesting stabilization of the triphasic rhythm, and decreased the probability of most other atypical states (Figure 9b, Table 3, *p < .*004, paired permutation test). Consistent with earlier studies that reported that RPCH can activate rhythms in silent preparations (Nusbaum and Marder, 1988), the probability of observing the silent state was driven to 0 in the presence of RPCH, together with other atypical states such as LP-silent and LP-silent-PD-bursting (Figure 9b).

**Table 3.** Probability distribution of states during modulator application, as shown in Figure 9

| State | Decentralized | RPCH | proctolin | oxotremorine | serotonin |
| --- | --- | --- | --- | --- | --- |
| regular | 0.39 | 0.73 | 0.69 | 0.78 | 0.27 |
| LP-silent | 0.06 | 0 | 0.02 | 0 | 0.07 |
| LP-silent-PD-bursting | 0.09 | 0 | 0.07 | 0 | 0.1 |
| PD-silent | 0.07 | 0 | 0 | 0 | 0.04 |
| PD-silent-LP-bursting | 0.01 | 0 | 0 | 0 | 0.03 |
| aberrant-spikes | 0.01 | 0.04 | 0.01 | 0.01 | 0.03 |
| LP-weak-skipped | 0.14 | 0.11 | 0.07 | 0.17 | 0.19 |
| PD-weak-skipped | 0.02 | 0.05 | 0 | 0 | 0 |
| sparse-irregular | 0.03 | 0 | 0.01 | 0 | 0.02 |
| irregular | 0.02 | 0.02 | 0.01 | 0 | 0.07 |
| silent | 0.07 | 0 | 0 | 0 | 0.01 |
| irregular-bursting | 0.1 | 0.04 | 0.11 | 0.03 | 0.17 |

**Table 4.** Code availability

| project | Notes |
| --- | --- |
| sg-s/crabsort | interactive toolbox to sort spikes from extracellular data |
| sg-s/stg-embedding | Contains all scripts used to generate every figure in this paper |
| KlugerLab/Flt-SNE | Fast interpolation based t-SNE, used to make embedding |
| sg-s/SeaSurfaceTemperature | wrapper to scrape NOAA databases |

**Figure 9.**
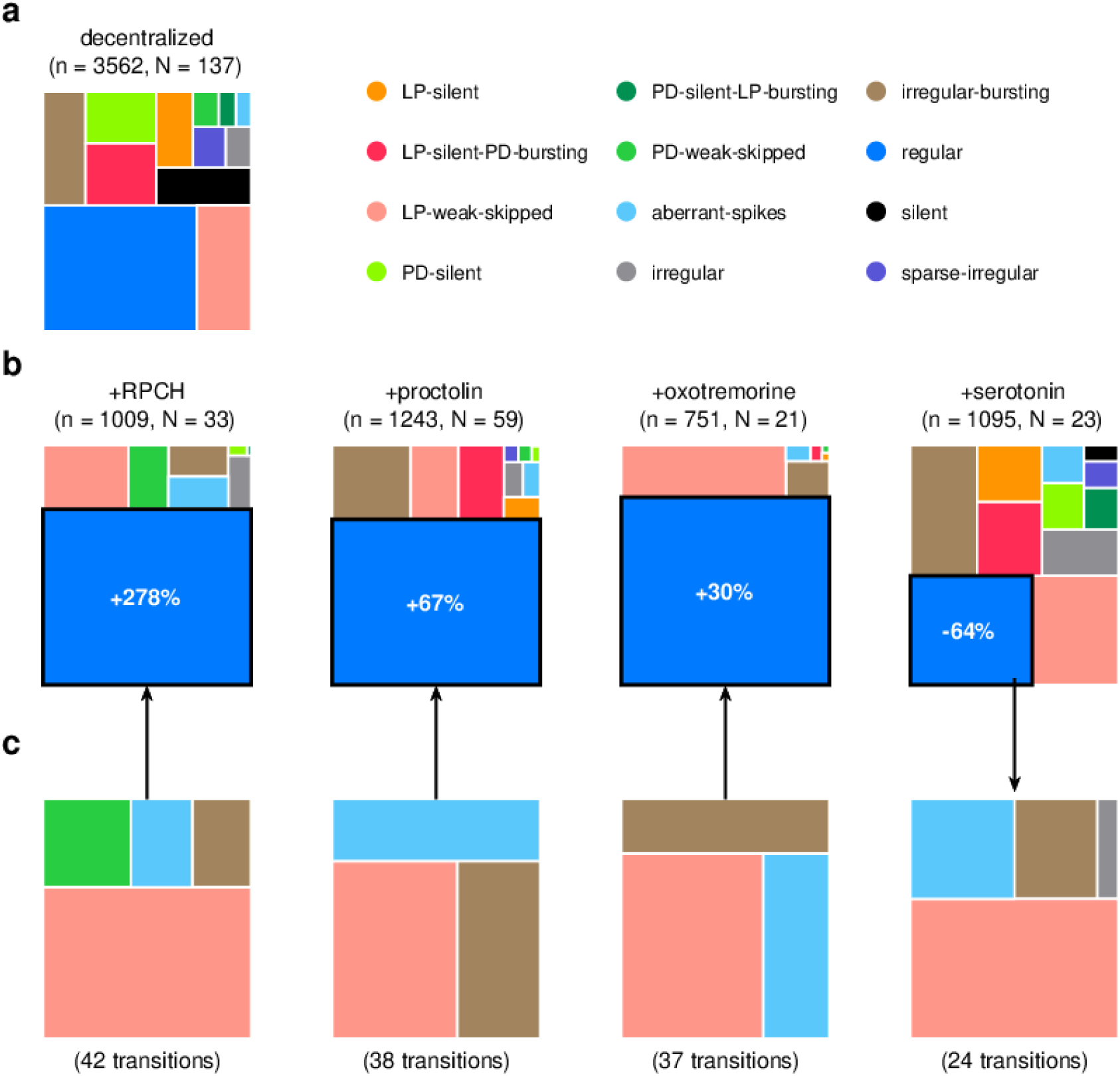
Effect of bath applied modulators. (a) State distribution in decentralized preparations. (b) State distribution on bath application of neuromodulators. Change percentages show difference in probability of regular state from decentralized to addition of neuromodulator. (c) Probability distribution of states conditional on transition to (for RPCH, proctolin and oxotremorine) or from (for serotonin) the regular state. (d) Coefficient of variation (CV) of burst periods of PD (purple) and LP (red) neurons vs. time before a transition away from `regular` states. *p, p* from Spearman test. *n* is the number of data points, *N* is the number of animals. **Figure 9-Figure supplement 1.** Raw traces during proctolin application **Figure 9-Figure supplement 2.** Neuromodulators affect map occupancy

Proctolin also targets a number of cells in the circuit (Swensen and Marder, 2001) and strengthens the pyloric rhythm through various mechanisms: by increasing the amplitude of slow oscillations in AB and LP (Hooper and Marder, 1987; Nusbaum and Marder, 1989), depolarizing the LP neuron (Golowasch and Marder, 1992; Turrigiano and Marder, 1993), and increasing the number of spikes per burst in LP and PD (Hooper and Marder, 1987; Marder et al., 1986; Hooper and Marder, 1984). Oxotremorine, a muscarinic agonist, has also been shown to enhance the robustness of the pyloric rhythm (Bal et al., 1994; Haddad and Marder, 2018; Rosenbaum and Marder, 2018). Similar to RPCH, both proctolin and oxotremorine significantly increase the probability of seeing the regular state (Figure 9b, Table 3, *p < .*004, paired permutation test), and the regular state is the only one significantly more likely when the neuromodulator is added. The strengthening effects of RPCH and oxotremorine are also manifested in the significantly lower probabilities of observing atypical and dysfunctional states such as silent, LP-silent, PD-silent, and sparse-irregular (Table 3).

Serotonin can have variable effects on the pyloric circuit, varying from animal to animal, and can either speed up or slow down the rhythm (Beltz et al., 1984; Spitzer et al., 2008). In *Panularis*, serotonin depolarizes LP in culture, but hyperpolarizes LP *in situ*, unlike other neuromodulators which typically have the same effect *in situ* and in culture (Turrigiano and Marder, 1993). Consistent with earlier work in *C. borealis* showing that serotonin destabilizes the rhythm in decentralized preparations (Haddad and Marder, 2018), we found that the probability of seeing `regular` states was significantly lower on addition of serotonin (Figure 9b, Table 3, *p < .*004, paired permutation test), together with a significantly higher probability of seeing atypical dysfunctional states such as `LP-silent, aberrant-spikes, PD-silent-LP-bursting` and `irregular`, suggesting loss of coordination between the many neurons in the pyloric circuit with serotonin receptors (Clark, 2004).

Do these modulators share common features in how they (de)stabilize the rhythm? We computed the probability distribution of states conditional on transitions to the regular state for RPCH, proctolin and oxotremorine, and conditional on transitions from the regular state for serotonin Figure 9c). For all four neuromodulators, the conditional state distribution was predominantly comprised of these three states: `LP-weak-skipped, irregular-bursting` and `aberrant-spikes`, suggesting that trajectories of recovery or destabilization of the regular rhythm share common features. Serotonin destabilizes the rhythm, decreasing the likelihood of observing `regular` states, similar to environmental perturbations (Figure 5) and decentralization (Figure 7).

Different neuromodulators activate different forms of the rhythm (Marder and Weimann, 1992; Marder and Hooper, 1985; Marder, 2012), partly because different neuron types express different receptors to varying extents (Garcia et al., 2015). Moreover, similar rhythmic motor patterns can be produced by qualitatively different mechanisms, such as one that depends on voltage gated sodium channel activity, and one that can persist in their absence (Harris-Warrick and Flamm, 1987; Epstein and Marder, 1990; Rosenbaum and Marder, 2018). To determine if different neuromodulators elicit regular rhythms that occupy different parts of the map, we plotted the location of data elicited by various neuromodulators in the full map (Figure 9–Figure Supplement 2). regular data elicited by different neuromodulators tended to lie in clusters, whose distribution in the map was significantly different between serotonin and CCAP, and proctolin and every other neuromodulator tested (*p < .*05, two-dimensional Kolmogorov Smirnoff test, using the method of Peacock (1983)). The differential clustering of `regular` states in the map with neuromodulator suggests that neuromodulators can elicit characteristic, distinct rhythms.

### Neuromodulators differentially affect transition between states

RPCH, proctolin and oxotremorine activate a common voltage dependent modulatory current, *I_MI_* (Swensen and Marder, 2001), but can differentially affect neurons in the STG because different cell types express receptors to these modulators to different degrees. For example, RPCH activates *I_MI_* strongly in LP neurons, but the effects of oxotremorine and proctolin are more broadly observed in the circuit (Swensen and Marder, 2000, 2001). Though these three modulators strengthen the rhythm, only rhythms elicited by oxotremorine and RPCH persist in tetrodotoxin, and proctolin rhythms do not, hinting that qualitatively different mechanisms underlie the generation of these seemingly similar rhythms (Rosenbaum and Marder, 2018). We therefore measured the transition rates between states during neuromodulator application to how similar or different trajectories towards recovery were.

In RPCH, proctolin and oxotremorine application, ≈ 100 transitions were observed between states (Figure 10). Transitions could not always be predicted by a null model assuming that transition probabilities scaled with the conditional probability of observing states after a transition. For example, some transitions, such as the transition from irregular to regular were never observed in RPCH, a significant deviation from the expected number of transitions given the likelihood of observing `regular` states after transitions (Methods and Materials). Others, such as the transition LP-silent to LP-silent-PD-bursting in proctolin and oxotremorine, were observed at rates significantly higher than expected from the null model. Strikingly, no transition is significantly over- or under-represented except the transitions from regular to `irregular-bursting` and to LP-weak-skipped across all three stabilizing modulators. Transitions into regular state are distributed across aberrant-spikes, LP-weak-skipped and `irregular-bursting` states for all three, but no invariant feature emerges in the rest of the transition matrix.

**Figure 10.**
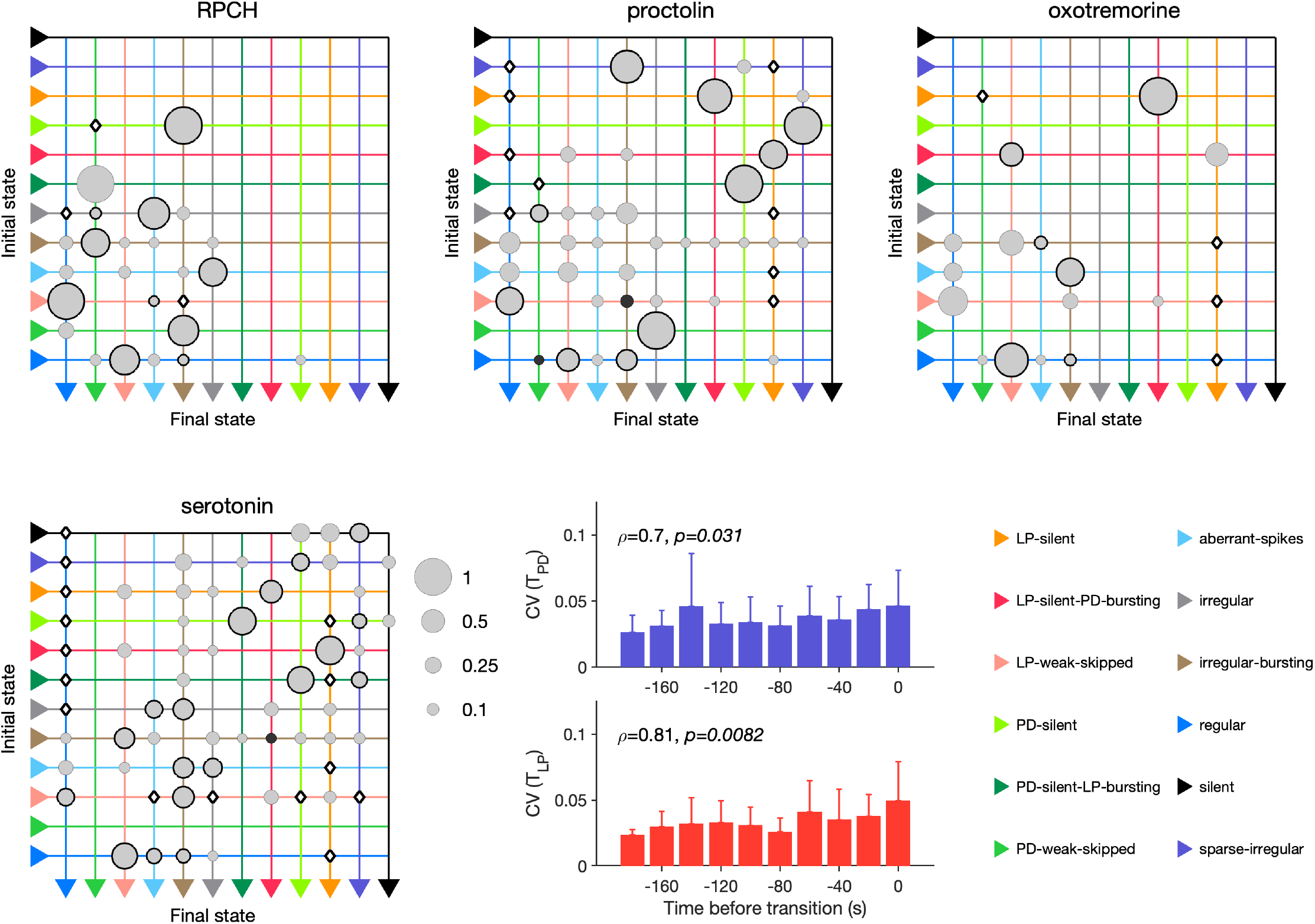
Effect of RPCH, proctolin, oxotremorine and serotonin on transition probabilities. Each matrix shows the conditional probability of observing the final state in the next time step given an observation of the initial state during bath application of that neuromodulator. Probabilities in each row sum to 1. Size of disc scales with probability. Discs with dark borders are transitions that are significantly more likely than the null model (Methods and Materials). Dark solid discs are transitions with non-zero probability that are significantly less likely than in the null model. O are transitions that are never observed, and are significantly less likely than in the null model. States are ordered from regular to silent. Bar graphics show the coefficient of variability (CV) of PD and LP burst periods before transition away from `regular` states. *p, p* from Spearman rank correlation test. RPCH: *n* = 148 transitions in *N* = 33 animals. Proctolin: *n* = 155 transitions in *N* = 59 animals. Oxotremorine: *n* = 102 transitions in *N* = 21 animals. Serotonin: *n* = 263 transitions in *N* = 23 animals. Bar graphs show the coefficient of variability (CV) of burst periods of PD and LP vs time before a transition away from `regular` states during serotonin application. *ρ, p* from Spearman rank correlation test.

Serotonin destabilizes the rhythm in decentralized preparations, and the transition matrix under serotonin reveals several features of the irregularity behavior observed under serotonin (Figure 10). A number of irregular and low-firing states states from `silent` to `irregular` never transition into the regular state, which is unlikely in the null model (*p < .*05, Methods and Materials). Transitions between pairs of states are symmetric and occur at rates significantly larger than in the null model, such as between LP-silent and LP-silent-PD-bursting. Intriguingly, destabilizing transitions from regular to LP-weak-skipped, aberrant-spikes and `irregular-bursting` are observed at rates signficantly higher than in the null model. These three abnormal states are also observed immediately preceding `regular` states in RPCH, proctolin and oxotremorine (Figure 9c), suggesting that mechanisms for both stabilization and destabilization of the rhythm share stereotyped trajectories.

Are transitions away from `regular` states also associated with increases in variability of burst periods? Similar to preparations in high [*K*^+^] and low pH, and when decentralized, transitions away from `regular` states in serotonin were associated with significantly rising variability in the burst periods of PD and LP neurons (Figure 10, *p < .*05, Spearman rank correlation test).

## Discussion

This study provides a concrete example of why it can be difficult to characterize experimental observations without the appropriate vocabulary to do so. Both highly stereotyped rhythms such as the pyloric oscillation, and highly irregular Poisson-like firing in large brain circuits are routinely described quantitatively using summary statistics. In the intermediate region between order and disorder, dynamics are harder to describe, and therefore frustrate efforts to systematically study circuits that generate them. We show that an unsupervised dimensionality reduction algorithm like t-SNE can create a useful representation of a dataset that is too large to visualize in its entirety using traditional methods. We incorporated domain-specific expert knowledge into this unsupervised approach by manually segmenting and labelling clusters in the embedding, identifying clusters of dynamics with biologically significant behavior. This dual approach conferred a two-fold advantage: both to more accurately measure traditional metrics such as burst metrics in `regular` states in large datasets (Figure 4, Figure 8), and to analyze irregular dynamics beyond the remit of conventional analysis methods (e.g., Figure 9). The map created in the present study (Figure 2) can be used as a blueprint to contextualize new experimental data from future experiments, which in turn can be added to the map to create a more complete picture of pyloric circuit dynamics.

### Robust identification of regular rhythms allows for detailed, interpretable analysis of rhythm metrics

Measuring the mean and variability of a regular oscillation in a neural circuit has several subtle challenges. Typically, variations in estimated metrics arising from cycle-to-cycle fluctuations are not distinguished from those arising from alteration between regions of regular bursting interrupted by regions of irregular spiking where these metrics are not defined. One way to disambiguate the two is to construct elaborate checks to make sure that the spike pattern being measured meets certain criteria. However, edge cases abound, and this is a challenging and poorly-motivated approach. One consequence of the embedding method we used is to reliably identify when rhythms were regular, and we found that burst metrics were well defined for this subset of data. We were therefore able to measure the mean and variability of various burst metrics (Figure 4), confident that we were measuring these metrics only in stretches of data where it made sense to do so. A byproduct of this restriction is that the variability in burst metrics measured this way stems almost entirely from cycle-to-cycle variations.

Consistent with years of study (Bucher et al., 2005; Hamood and Marder, 2015; Hamood et al., 2015), our results (Figure 4) show explicitly that within-animal variability in pyloric burst metrics is less than across-animal variability. Our results are from a meta-analysis of data from several different experimenters from different laboratories, collected over a span of ten years. It is therefore an ideal dataset in which to measure variability. We find that the coefficient of variation of all burst metrics measured is ≈ 0.1 (Figure 4b), which is a proxy for how regular the pyloric oscillation can be under baseline conditions. Measuring burst metrics on decentralization (Figure 8) also allowed us to characterize how regular rhythms change, while still being recognizably regular. In addition to recapitulating well-understood phenomena such as the slowing down and increased variability in rhythms, we found that phase of LP burst starts did not significantly change, but phases of LP bursts stops did, suggesting that features of the rhythm are differentially robust to the removal of neuromodulation.

### Numerical methods to analyze neural circuit dynamics

Advances in experimental techniques in neuroscience allow for recordings from larger number of neurons for longer periods. There have been contemporaneous advances in techniques to analyze this data. A first step in data analysis is often data visualization. Modern neural data can be large and high dimensional, and visualizing the entirety of a large data set can be a non-trivial task. Visualization and other forms of data analysis rely on dimensionality reduction (Nguyen and Holmes, 20fi9).

Here we used the t-SNE algorithm as a core method to reduce the dimensionality of the dataset and to visualize our data. t-SNE has been widely used in the unsupervised analysis of many types of biological data (Berman et al., 2014; Kollmorgen et al., 2020; Chen et al., 2020; Macosko et al., 2015; Kobak and Berens, 2019; Leelatian et al., 2020), including neural recordings (Dimitriadis et al., 2018). t-SNE minimizes the Kullback-Leibler divergence between a Gaussian distribution modeling pairwise distances between data points and a Student t-distribution modeling distances between the same points in a low (typically two) dimensional embedding (Van der Maaten and Hinton, 2008; Linderman and Steinerberger, 2019). This feature makes t-SNE an attractive tool to try to visualize data sets such as the data in this paper, because it can demonstrate how similar spike patterns are to each other.

t-SNE has also been used to find clusters in data, since its original use in visualizing and clustering hand-written digits in the MNIST database (Van der Maaten and Hinton, 2008). t-SNE has been shown rigorously to be capable of recovering well-separated clusters (Linderman and Steinerberger, 2019). Neighborhood embedding techniques like t-SNE combine attractive forces between pairs of points with repulsive forces between all points. Stronger attraction can better represent smoothly varying manifold structures, while stronger repulsion can better represent discrete cluster structures (Böhm et al., 2020). In our application, t-SNE generated clusters where spike patterns could be described as qualitatively different. For example, spike patterns in top-most cluster (colored green in Figure 3) all had weak PD spiking, but regular and strong LP spiking. This was qualitatively different from the two closest clusters LP-weak-skipped and irregular. In regions of the map where clusters were not cleanly separated (for example, in the connection between the regular and `irregular-bursting` clusters), manual inspection revealed a number of intermediate states. The “clustered” or “not-clustered” regions of the map are therefore informative of the underlying distribution of spike patterns, and emerge robustly from the embedding.

t-SNE-based methods are not the only way to analyze such data, and a variety of other methods have been developed recently. Multidimensional Scaling (MDS) (Cox and Cox, 2008) has been used to visualize collective coding for different task dimensions in a population of neurons in the amygdala in rats (Kyriazi et al., 2018). Convolutional non-negative matrix factorization (Mackevicius et al., 2019) has been used to find sequences in neural and behavioral data by building a parts-based representation of the data. Recent work (Williams et al., 2020) extends this method by including a point process model to model sparse spike sequences without binning time. Tensor Component Analysis (Williams et al., 2018) can generate three low-dimensional descriptions from neural data: separating out neuron-specific, trial-specific and temporal factors, making it valuable in multi-trial data. Dynamical Component Analysis is a linear method that attempts to find dynamics rather than explaining variance in the data (as in PCA) (Clark et al., 2019).

Methods based on neural networks offer powerful tools to analyze unstructured neural data. Generally, one method to study how a high-dimensional neural system works is to model it with a recurrent neural net (RNN), and then to study the RNN model (Vyas et al., 2020). Autoencoders offer an interesting way of dimensionality reduction (or latent space analysis) because their architecture contains an information bottleneck (Rumelhart et al., 1985), and have long been a focus of unsupervised machine learning (Baldi, 2012). Topological autoencoders combine autoencoders with the concept of persistent homology, and use a topological loss term that minimizes differences between the topological signatures of the data and the representation in the lower dimensional space (Moor et al., 2019). These methods are similar in spirit to the analysis presented here, but use sophisticated neural nets whose parameters yield the lower-dimensional representation. Other end-to-end analysis methods include a method called SOM-VAE, which combine self-organizing maps (SOMs) and variational auto-encoders (VAEs)(Fortuin et al., 2018) to analyze high dimensional time series and find transitions between states, and deep temporal clustering, which combines dimensionality reduction and temporal clustering in a single unsupervised learning problem (Madiraju et al., 2018).

### Applicability to bigger circuits and unidentified neurons

In this study, we used spiking patterns of PD and LP cells as a proxy for the dynamics of the pyloric circuit. A better characterization of pyloric circuit dynamics would include AB, PY, VD and IC cells (Eisen and Marder, 1982; Marder and Bucher, 2007). The data analyzed in this study did not consistently have recordings that made it possible to reliably and consistently extract spike times of the VD and PY neurons. While most of the data included recordings from the *lvn* nerve, extracting PY spike times from the *lvn* was not feasible at scale. Spikes from PY are smaller than spikes from LP and PD on the *lvn*, and the duration of PY bursting may partially overlap with that of LP and PD. Even when data include recordings from the *pyn*, identifying PY spikes is not straightforward. There are several PY neurons, whose spikes may overlap to varying degrees to variability in the subpopulation of PY neurons that spike and the precise timing of action potential initiation. The shape of PY spikes can therefore be quite variable. In addition, spikes from the gastro-pyloric LPG neuron are often observed on the *pyn* (Figure 1b). Even intracellular recordings from PY neurons are not necessarily sufficient to accurately estimate PY spike times because intracellular recordings measure the activity of only the cell recorded from, and it is not uncommon to observe that the PY cell being recorded from generates fewer spikes than the other PY neurons as observed extracellularly, possibly due to leak currents introduced from sharp electrodes (Cymbalyuk et al., 2002).

In this analysis, we chose to include features such as the “spike phase” (Figure 2b-c) because the neurons in this circuit are mutually coupled with inhibitory and electrical synapses and therefore strongly affect the activities of each other in the collective rhythm. An analysis of circuit dynamics from other neural networks that did not show such strong intrinsically phase-controlled behavior could use other features more suitable to those systems. The analysis method in this study is well-suited for large datasets of neural recordings from identified neurons. Data where the identity of each neuron is not uncontrolled, or cannot be known, such as large scale recordings from a brain, would require modifications to the analysis pipeline described in Figure 2. First, it would no longer be possible to construct a data vector of fixed length, because ordering of the different neurons would not be meaningful. Each data point would instead be an unordered set of spike times from each neuron, and a distance function that operated on spike times (Christen et al., 2006; Victor and Purpura, 1997; Schreiber et al., 2003; Rossum, 2001) could be used to generate a distance matrix between raw data points, which would be the input to the embedding algorithm.

### Ahead-of-time experimental design can maximize utility and interpretability of data

This study used a large dataset collected by various experimenters, and included data originally collected for other studies (Tang et al., 2012, 2010; Haddad and Marder, 2018; Haley et al., 2018; Rosenbaum and Marder, 2018; Powell et al., 2021; He et al., 2020). As such, this post-hoc analysis is limited ultimately by the data: its quantity, the way it was collected, and the decisions made and tradeoffs chosen by the experimenter who collected it. A general lesson learned here is that close coordination between experimenters and theorists and data analysts can help maximize the utility of data collected. Because experiments are expensive to perform, in the time of researchers, reagents and experimental animals, seemingly inconsequential changes to the way data are collected can substantially increase the amount of usable data to a greater number of questions, some of which may not be well-formulated at the time of data collection.

For example, studying the effects of perturbations to pyloric circuits forces experimenters to make choices about experimental protocol that have far-reaching consequences on the analysis and interpretation of data collected. If perturbations are severe enough to destabilize the pyloric rhythm, and even cause prolonged periods of silence, should an identical sequence of perturbations be used in every preparation if some preparations “crash” under relatively moderate perturbations and greater perturbations may risk irreversible changes? Is it more important to introduce perturbations that change at a certain, fixed rate, or should perturbation intensity be dialed up or down based on observed responses, to better characterize the full response range of the system being perturbed? Experimental constraints and the priorities of specific studies have led to a patchwork of choices in the dataset used here, which means that it is not entirely straightforward to disentangle the effects of applied perturbations at a given time from the cumulative effects of the entire experimental protocol and stimulus history.

Data acquisition systems allow experimenters to record from neurons at high temporal resolution for long periods of time. Time-varying metadata, such as pH, temperature, [*K*^+^] or the concentration of added modulators are not always recorded concomitantly, because they can be difficult to measure. Temperature and pH probes, when used, can yield high-resolution and automatic logging of these quantities. Because decentralization involves a manual intervention such as cutting the *stn* or constructing and filling a well on the *stn*, and because the process takes time, the precise time of decentralization can be hard to record and estimate, leading to a fraction of preparations being decentralized before the nominal start of decentralization, with effects being evident as in the apparent increase in burst period shown in Figure 8d.

### Cryptic circuit variability can be revealed by diversity in crashes

A large body of work has shown that there is more than one way to make a functional neural circuit (Prinz et al., 2003, 2004; Gutierrez et al., 2013). Several combinations of circuit parameters such as synapse strengths, ion channel conductances and network topology can be found in circuits that generate similar emergent collective dynamics (Gonçalves et al., 2020). In the pyloric circuit, the dimensionality of the space of neuron and circuit parameters is larger than the dimensionality of the rhythm: ≈ 50 parameters are required to specify ionic and synaptic conductances even in simplified models, but the rhythm under baseline conditions can be well described using a handful of metrics (Marder and Bucher, 2007). This disparity in dimensionality leads to an inherently many-to-one mapping from the space of circuit architecture to the space of circuit dynamics. Pyloric circuits at baseline can therefore exhibit “cryptic” architectural variability (Haddad and Marder, 2018), where the diversity of circuit topologies and neuron parameters underlying functional circuits is masked by the relatively low-dimensional nature of the observation of regular rhythms. Intriguingly, there was no seasonal effect on the variations in bursting under baseline conditions (Figure 4–Figure Supplement 4), or sensitivity to decentralization (Figure 7–Figure Supplement 4), suggesting that these dimensions of observed variability my arise from other factors such as circuit-to-circuit architectural differences.

Perturbations can reveal differences between seemingly identical circuits because parameter differences that were inconsequential in the generation of baseline activity can now generate disparate dynamics. Perturbations such as current injections in a network of oscillators can shift phases, revealing connection weights between individual neurons (Timme, 2007). What do the perturbations used in this work do? Some perturbations like decentralization can have complex, time varying and variable effects, because neurons in the STG are multiply modulated (Marder, 2012); this may lead to the complex and diverse responses seen on decentralization (Figure 7). Others like changing extracellular [*K*^+^] can have more focussed effects, which changes the reversal potential of *K*^+^ ions, altering currents through *K*^+^ permeable ion channels, and tends to depolarize neurons (He et al., 2020). The challenge in interpreting data from experiments with perturbations such as these is the dual complexity of the elicited circuit behavior and the functional effects of the perturbations. Future work with other, sparse perturbations can help determine if diverse dynamics observed in the present work are a consequence of the complex nature of the perturbations used.

If there are many solutions to a designing a functional circuit, are some solutions more robust to all perturbations? Alternatively, is there a tradeoff for circuits between being robust to perturbation *x* and being robust to perturbation *Y*At the population level, some animals could possess pyloric circuits more robust to one perturbation, and the expense of greater sensitivity to another; and other animals could possess circuits that are more robust to other perturbations. Recent work studying a population of isolated pacemaker kernels of the pyloric circuit (AB and PD cells) found only moderate correlation between robustness to perturbations in pH and temperature (Ratliff et al., 2021). Examples of population-level hedges against uncertain environmental perturbations include the diversity in chemotactic behavior in bacteria (Frankel et al., 2014). In modeling work with neurons, recent work has shown that homeostatic regulation rules that confer robustness to some perturbations can create sensitivity to other perturbations (O’Leary et al., 2014; Gorur-Shandilya et al., 2020).

### Linking behavior to mechanisms

The present work offers a path towards analysis that can reveal cryptic variability and build mechanistic links from circuit architecture to function. By characterizing the totality of circuit dynamics under a variety of conditions, this study equips further work with the tools to fit biophysically detailed models of the pyloric circuit to diverse circuit dynamics under baseline conditions and perturbations. From the large diversity of neuron and circuit parameters that can reproduce a snapshot of activity, will only a subset of models recapitulate the diverse irregular behavior seen under extreme perturbations? Recent work that reproduced how circuits change cycle periods with temperature (Alonso and Marder, 2020) can be extended to find parameter sets that also generate the irregular states characterized in this study, at the rates observed in the data, and will help resolve this question. Future experimental work can pair data analysis methods such as this work with quantitative measurements of cellular and circuit parameters using emerging techniques (Schulz et al., 2006, 2007; Tobin et al., 2009) to find parameter values of cells that generate robust rhythms and irregular states.

### Diversity and stereotypy in trajectories from functional to crash states

Are there preferred paths to go from regular rhythms to crash? Diversity in the solution space of functional circuits, and the varied effects of perturbations on these circuits, argue for an assortment of trajectories from function dynamics to irregular or silent states. While transition matrices measured during different perturbations were varied (Figure 6), we did observe universal features in transition matrices measured during environmental perturbations, decentralization, and addition of neuromodulators (Figure 6, Figure 7, Figure 10). The destabilizing transition from regular --- LP-weak-skipped was over-represented in every transition matrix, suggesting that the weakening of the LP neuron is a crucial step in the trajectories towards destabilization, perhaps because there is only one copy of LP in the circuit. Earlier work studying trajectories of destabilization of regular bursting in the isolated pacemaker kernel also found a common trajectory of destabilization, from regular bursting to tonic spiking to silence (Ratliff et al., 2021). Transitions away from regular rhythms were also associated with increased variability in burst periods during all perturbations except high temperature and low pH (Figure 6, Figure 7, Figure 10). Earlier work on the isolated pacemaker kernel found similar increase in variability in PD voltage dynamics before transitions from regular bursting, similar to the increasing variability measured in the present study (Ratliff et al., 2021).

The structure of the transitions between states also hints at features of the circuit that are critical for rhythm (de)stabilization. Unsurprisingly, PD-silent states precede silent states in low pH, high temperature and high [*K*^+^] perturbations (Figure 6). This makes sense because PD cells are electrically coupled to the endogenous burster AB in the pacemaker kernel, and silencing the pacemaker kernel can cause the circuit to go silent. Though the states are determined purely from clusters in the embedding (Figure 2), and thus from statistical features of spike times, some states may be identified predominantly with cell-specific features (e.g., LP-weak-skipped where the LP neuron fails to burst regularly, but the PD neurons do), or with circuit-level features (e.g., aberrant-spikes where one or both neurons fire spikes outside the main burst, which may be caused by incomplete inhibition from the reciprocal neuron). Decentralization elicits the largest number of transition types, with ≈ 80% of all transition types observed, which could be a consequence of the complex change in the neuromodulator milieu following transection of descending nerves.

### Comparison with other categorization methods

Earlier work categorized the varied dynamics of the pyloric circuit during perturbations (Haddad and Marder, 2018; Haley et al., 2018; Ratliff et al., 2021; Alonso and Marder, 2020). In that work, categories were typically constructed by hand and were not rigorously shown to be mutually exclusive. Categories in the present work, while being manually chosen, emerge from the distribution of the data in the map (Figure 3); and no segment of data can have more than one label, because it can exist only at a single point in the map. Earlier work categorized rhythms that were labelled regular into two categories, “normal triphasic” and “normal triphasic slow” (Haddad and Marder, 2018). While there is significant variation in the burst periods in the regular cluster, (Figure 2–Figure Supplement 1, Figure 4), we did not observe a distinctly bimodal distribution of burst periods, and therefore could not justify splitting regular into two. Earlier work also included a category called “gastric like rhythms”, where LG or DG neurons were active, indicating the presence of the gastric mill (Weimann and Marder, 1994). Because the present work only considers spikes on PD or LP neurons, circuit dynamics with gastric activity are scattered across states, based on how the gastric activity affects PD and LP spikes. The “LP01” state identified in Haddad and Marder (2018) is equivalent to the LP-weak-skipped state; the present work also identified a PD-weak-skipped which was not identified in the earlier work, perhaps because it is ≈ 1/7 as prevalent (Figure 3). The catch-all “atypical firing” state could be teased apart into a number of irregular states (irregular, `irregular-bursting`, sparse-irregular) that span several well-separated clusters in the map (Figure 3). In summary, the present work recapitulates every label constructed to categorize spike patterns from PD and LP neurons in earlier work, and additionally finds new spike patterns that were either not detected or not identified as distinct.

Our work provides a key tool to characterize non-regular spike patterns in small neural circuits and thus provides a bridge between experimental or simulation work grounded in the biophysical detail of ion channels and synaptic currents; and the rich body of observations of circuits under baseline and challenging conditions. The tools we have employed can easily be adapted to other circuits and systems, makes limited assumptions of the dynamics of the circuit, yet provides a robust framework on which to hang a large volume of previously ineffable expert domain knowledge.

## Methods and Materials

### Animals and experimental methods

Adult male Jonah crabs (*C. borealis*) were obtained from Commercial Lobster (Boston, MA), Seabra’s Market (Newark, NJ) and Garden Farm Market (Newark, NJ). Dissections were carried out as previously described (Gutierrez and Grashow, 2009). Decentralization was carried out either by cutting the *stn*, or by additionally constructing a well on the *stn* and adding sucrose and TTX as described in Haddad and Marder (2018). Temperature was controlled as described in Tang et al. (2012, 2010); Haddad and Marder (2018). Extracellular potassium concentrations were varied as described in He et al. (2020). pH perturbations are described in Haley et al. (2018).

### Spike identification and sorting

Spikes are identified from extracellular recordings of motor nerves or from intracellular recordings. LP spikes were identified from intracellular recordings, *lvn*, *lpn* and *gpn* nerves (in descending order of likelihood). PD spikes were identified from *pdn*, intracellular recordings, and *lvn*. We used a custom-designed spike identification and sorting software (called “crabsort”) that we have made freely available at https://github.com/sg-s/crabsort previously described in Powell et al. (2021). Spikes are identified using a fully connected neural network that learns spike shapes from small labelled data sets. A new network is typically initialized for every preparation. Predictions from the neural network also indicate the confidence of the network in these predictions, and uncertain predictions are inspected and labelled and the neural network learns from these using an active learning framework (Settles, 2009).

### Data curation and data model

Each file was split into 20-second non-overlapping bins and spike times, together with metadata, were assembled into a single immutable instance of a custom-built class (embedding.DataStore). The data store had the following attributes:

• *spike times* containing LP and PD spike times.

• *ISIs* containing inter-spike intervals and spike phases

• *labels* categorical data containing manually generated labels from Figure 3

• *metadata* such as concentration of modulators, pH, temperature, whether the preparation was decentralized or not, etc.

Using an immutable data structure reduced risks of accidental data alteration during analysis. Every attribute was defined for every data point.

### Embedding

#### ISI and phase representation (*Figure 2*b)

Each data point is a 20-second bin containing spike times from LP and PD neurons (Figure 2a) . For each data point, spike times are converted into inter-spike intervals. A set of spike times uniquely identifies a set of (ordered) inter-spike intervals. The set of LP spike times generates a set of LP ISIs, and the set of PD spike times generates a set of PD ISIs (Figure 2b).

For every spike in PD or LP, a “spike phase” can be calculated as follows. Spike phases are not defined when either LP or PD are silent in that data point, or for LP/PD spikes with no spikes from the other neuron before or after that spike. Thus the “spike phase” of the *i*-th spike on neuron X w.r.t neuron Y is given by:

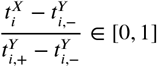

where 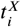 is the time of the *i*-th spike on neuron 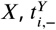 is the time of the last spike on *Y* before 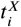 and 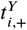 is the time of the first spike after 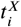. Note that this definition can be generalized to *N* neurons, though the number of spike phases grows combinatorially with *N*.

#### Construction of vectorized data frame (*Figure 2*c-d)

Each data point can contain an arbitrary number of spikes, and thus an arbitrary number of ISIs and spike phases. Ideally, each data point is a data frame of fixed length (a point in some fixed high-dimensional space). To do so, we computed percentiles ISIs and spike phases (Figure 2c). We chose ten bins per ISI type (deciles). The end result is not strongly dependent on the number of bins chosen, as long as there are sufficiently many bins to capture the distinctly bimodal distribution in ISIs during bursting.

We included three other features to help separate spike patterns that appeared qualitatively different. First, firing rates of LP and PD neurons. Second, the ratios of 2nd-order to fist-order ISIs, defined as:

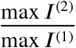

where *I*^(^*^n^*^)^ is the *n*-th order set of ISIs computed as time the time between *n* spikes. *I*^(1)^ is the simple set of ISIs defined between subsequent spikes. This measure is included because it captures the difference between single spike bursts and normal bursts well.

Finally, we also included a metric defined as follows:

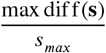

where **s** is a vector of sorted ISIs and *s_max_* is the sorted ISI for which the difference between it and the previous sorted ISI is maximum. This metric was included as it captures to a first approximation how “burst-like” a spike train is. Intuitively, this metric is high for spike trains with bimodal ISI distributions, as is the case during bursts.

All these features were combined into a single data frame and *z*-scored (Figure 2d).

In some cases, these features were not defined, e.g.: when there are no spikes on either neuron, the concepts of spike phases or ISIs are meaningless. In these cases, “filler” values were used that were located well off the extremes of the distribution of the metric when defined. For example, ISIs were filled with values of 20s (the size of the bin) when no spikes were observed. The overall results and shape of the embedding did not depend sensitively on the value of the filler values used.

#### Embedding using t-SNE

So far, we have described how we converted a 20-second snippet containing spike times from LP and PD into a data frame (a vector). We did this for every 20-second snippet in the dataset. Data that did not fit into any bin was discarded (for example, data at the trailing end of an experiment shorter than 20 seconds). Thus, our entire dataset is represented by *M* × *N* matrix where *M* is the number of features in the data frame and *N* is the number of data points.

We used the t-SNE algorithm (Van der Maaten and Hinton, 2008) to visualize the vectorized data matrix in two dimensions. Our dataset contained ≈ 10^5^ points, and was therefore too large for easy use of the original t-SNE algorithm. We used the FI-tSNE approximate algorithm (Linderman et al., 2019) to generate these embeddings. We used a perplexity of *P* = 100 to generate these embeddings. Varying perplexity caused the embedding to change in ways consistent with what is expected for t-SNE embeddings, and the coarse features of the embedding did not sensitively depend on this choice of perplexity (Figure 2–Figure Supplement 3).

t-SNE is often used with random initialization, and different random initializations can lead to different embeddings with clusters located at different positions in the map. The importance of meaningful initializations has recently been highlighted (Kobak and Linderman, 2021), and we used a fixed initialization where the X-axis corresponded to the shortest ISI in each data point, and the Y-axis corresponded to the maximum ratio of 2nd-order to fist order ISI ratios (described above). For completeness, we also generated embeddings using other initializations (Figure 3–Figure Supplement 2). For both random initializations (Figure 3–Figure Supplement 2a-d) and initializations based on ISIs (Figure 3–Figure Supplement 2e-f), we observed that `regular` states tended to occur in a single region, surrounded by clusters that were dominated by a single color corresponding to irregular states. Thus, the precise location of different clusters can vary with the initialization, but the overall structure of the embedding, and the identity of points that tend to co-occur in a cluster, does not vary substantially with initialization.

#### Triangulation and triadic differences (*Figure 2–Figure Supplement 2*)

The output of the embedding algorithm is a set of points in two dimensions. We built a Delaunay triangulation on these points. For each triangle in the triangulation, we computed the maximum difference between some burst metric (e.g., burst period of PD neurons) across the three vertices of that triangle. These triadic differences are represented colored dots, where the dots are located at the incenters of each triangle in the triangulation.

### Time series analysis

#### Measuring transition matrices (*Figure 6, Figure 7, Figure 10*)

The transition matrix is a square matrix of size *N* that describes the probability of transitioning from one to another of *N* possible states. The transition matrix we report is the right stochastic matrix, where rows sum to 1. Each element of the matrix *T_ij_* corresponds to the conditional probability that we observe state *j* given state *i*. To compute this, we iterate over the the sequence of states and compare the current state to the state in the next state. Breakpoints in the sequence are identified by discontinuities in the timestamps of that sequence and are ignored. We then zeroed the diagonal of the matrix and normalized each row by the sum.

Measuring variability before transitions away from `regular` states (Figure 6, Figure 7) We first identified continuous segments that corresponded to uninterrupted recordings from the same preparation at the appropriate condition. For each segment, we found all transitions away from the regular state. We therefore computed a vector, as long as the segment, containing the time to the next transition. We then collected points corresponding to time to next transition ranging from *t* = −200*s* to *t* = 0*s*. For each time bin, we measured the coefficient of variation of the burst period by dividing the standard deviation of the burst period in that datum by the mean in that datum.

### Data visualization

#### Raincloud plots (*Figure 4*)

Raincloud plots (Allen et al., 2019) are used to visualize a univariate distribution. Individual points are plotted as dots and a shaded region indicates the overall shape of the distribution. This shape is obtained by estimating a kernel smoothing function estimate over the data. Individual points are randomly jittered along the vertical axis for visibility.

#### Occupancy maps (*Figure 5, Figure 7*)

To visualize where in the map data from a certain condition occurred, the full embedding is first plotted with colors corresponding to the state each point belongs to. The full dataset is made semi-transparent and plotted with larger dots to emphasize the data of interest. Data in the condition of interest is then plotted as usual. Each bright point in these plots corresponds to a 20-second snippet of data in the condition indicated.

#### Treemaps (*Figure 7, Figure 9*)

Treemaps (Shneiderman and Wattenberg, 2001) were used to visualize state probabilities in a given experimental condition. For each preparation, the probability of each state was computed, and the mean probability of a given state was computed by averaging across all preparations. Thus, each preparation contributes equally. The area of the region in the treemap scales with the probability of that state.

#### Transition matrices (*Figure 6, Figure 7, Figure 10*)

Transition matrices were visualized as in Corver et al. (2021). Initial states are shown along the left edge and final states are shown along the bottom edge of each matrix. Lines are colored by origin (horizontal lines) or destination (vertical) states. The size of each disc at the intersection of each line scales with the conditional probability of moving from the initial state to the final state. Note that the size of all discs is offset by a constant to make small discs visible.

### Statistics

#### Comparing within-group to across-group variability (*Figure 4*)

To compare the variability of various burst metrics within each animal and across animals, we first measured the means and coefficients of variations (CV) of each burst metrics in every animal. We then used the mean of the coefficients of variations as a proxy for the within-animal variability, and used the coefficient of variation of the means as a proxy for the across-animal variability. Note that both measures are dimensionless. They can therefore be directly compared.

To test if the within animal variability was significantly less than the across animal variability, we performed a permutation test. We shuffled the labels identifying the animal to which each data point belonged to and measured a new “within-animal” and “across-animal” variability measure using these shuffled labels. We repeated this process 1000 times to obtain a null distribution of differences between within- and across-animal variability. Identifying where in the null distribution the data occurred allowed us to estimate a *p*-value for the measured difference. For example, if the measured difference between within- and across-animal variability in metric X was greater than 99% of the null distribution obtained by shuffling labels, we conclude that the *p*-value is .01. The significance level of .05 was divided by the number of burst metrics we tested to determine if any one metric was significantly more or less variable across animals.

#### Comparing map occupancy before and after decentralization (*Figure 7*b)

To determine if data are more widely distributed in the map after decentralization, we computed the mean distances travelled in the map between subsequent time points for each preparation. Each preparation’s circuit dynamics is represented as a trajectory in this map. Distances in the map between subsequent points are measured and summed for each preparation.

Each point in (Figure 7b) corresponds to a single preparation before and after decentralization. Data are therefore paired and we can generate a null distribution by randomly shuffling each pair. This null distribution is shown in the gray shading in (Figure 7b). The dashed line is the line of unity and indicates the middle of the null distribution. The measured difference between the distances travelled in the decentralized and intact cases is shown in the purple line. The *p* value can be estimated as the fraction of the null distribution greater in magnitude than the purple line.

#### Measuring trends in variability in regular rhythms before transitions (*Figure 6*b *,Figure 7*f*, Figure 9*d)

To determine if variability significantly increased in the 200s preceding a transition away from regular, we measured the Spearman rank correlation between time before transition (x-axis) and mean variability. The Spearman rank correlation *ρ* is 1 if quantities monotonically increase.

#### Measuring transition rate significance (*Figure 6*a *,Figure 7*e*, Figure 10*)

In the empirical transition matrices, certain transitions never occur, and certain transitions occur with relatively high probability. Each element of the transition matrix *T_ij_* corresponds to the conditional probability *P* (final initial). Our null model assumes that transitions occur at random between states, and therefore the probability of observing any transition *i* --- *j* scales with the marginal probability of observing state *j* after transitions. We therefore built a null distribution of transition rates by sampling with replacement from the marginal counts of states after transitions. The fraction of this null distribution that was above or below the empirical transition rate was interpreted to be the *p*-value and thus determined significance.

### Code availability

The following table lists code used in this paper. Code can be downloaded by prefixing https://github.com/ to the project name.

## Acknowledgments

This paper includes data collected by Lamont Tang, Lily He, Mara Rue, Jessica Haley, Daniel Powell, Anatoly Rinberg, and Ekaterina Morozova. We gratefully acknowledge helpful conversations with Paul Miller, Mark Zielinksi, Sriram Sampath and Alec Hoyland.

This work was funded by NIH grants T32 NS007292 (SGS) and R35 NS097343 (EM and SGS), NIH MH060605 (FN and DB) and DFG SCHN 1594/1-1 (ACS).

---

**Figure 2–Figure supplement 1.**
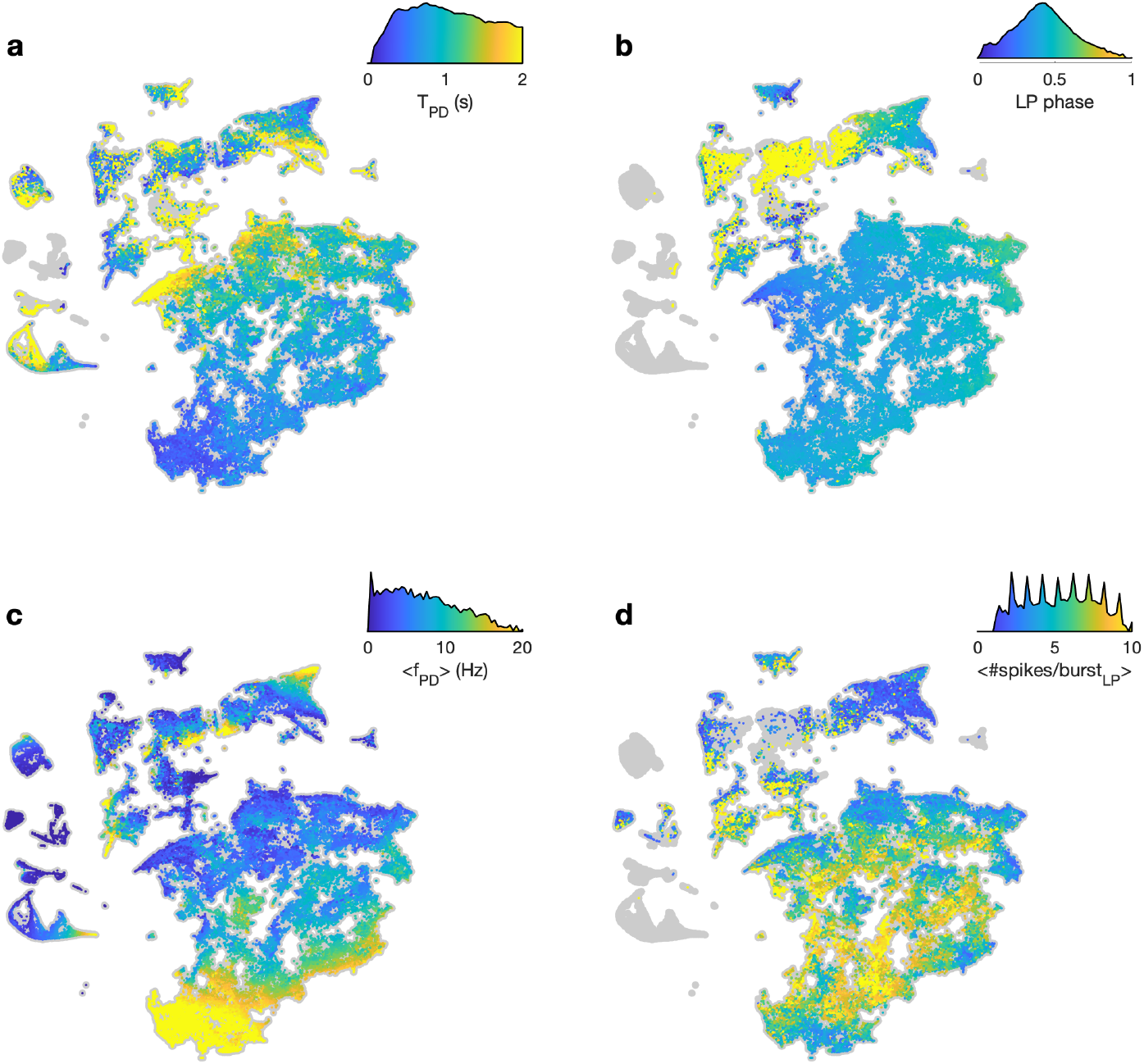
Burst metrics smoothly vary in map. In each panel, embedding of the entire dataset is shown in gray. Points are colored by (a) burst period of the PD neuron (b) phase of LP burst start in PD time (c) mean firing rate of PD neuron and (d) mean number of spikes per burst in the LP neuron. In each panel, the color scale also shows the distribution of metric over the entire data set (Y-axis in log scale). The distribution in (d) is spiky because the mean number of spikes/burst tends to be integer valued.

**Figure 2–Figure supplement 2.**
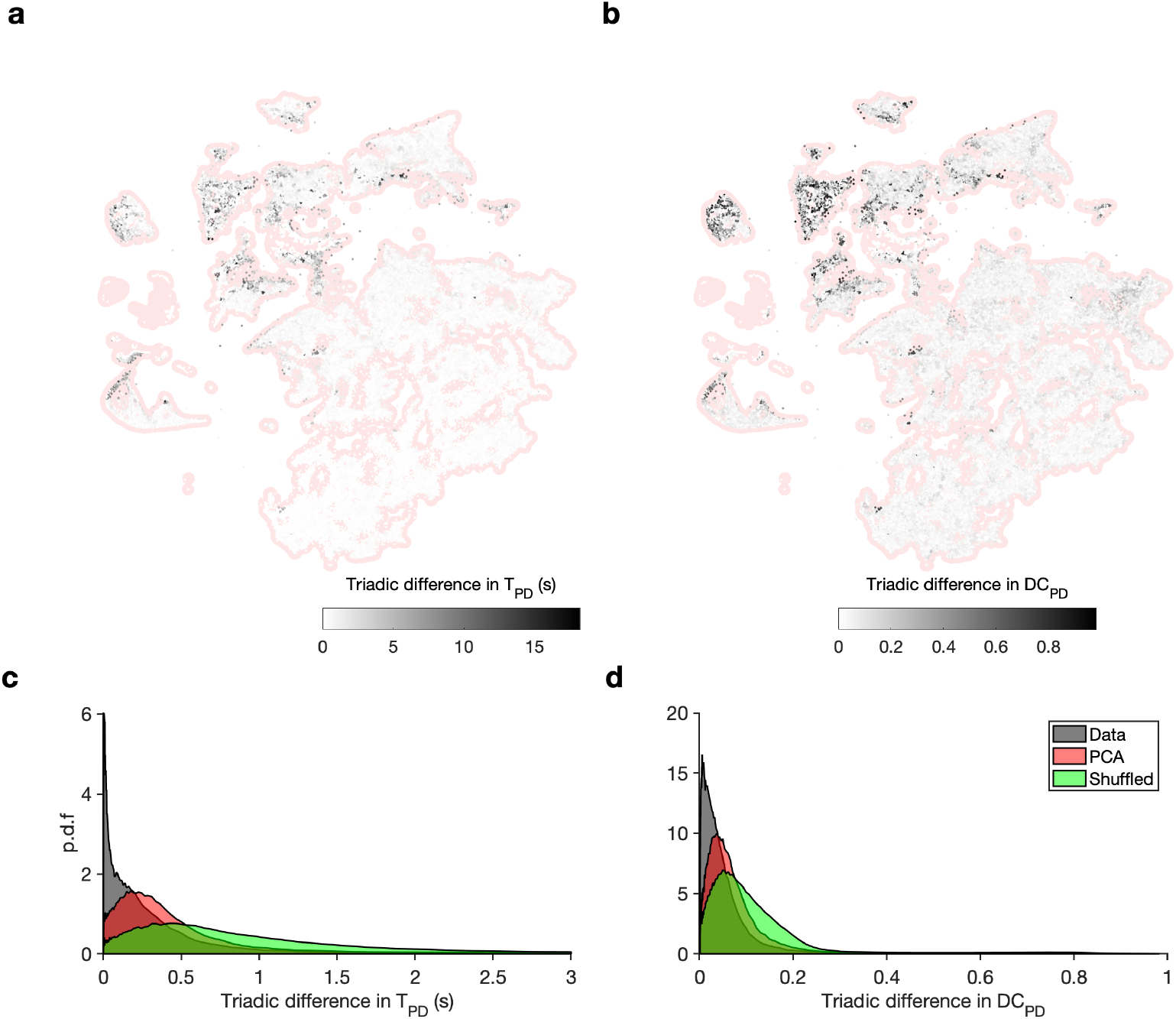
Embedding arranges data so that neighbors tend to be similar. (a) Map shows embedding of all data in red shading. Shaded dots are incenters of a Delaunay triangulation of the map, and shading indicates the maximum absolute value of difference between burst period of the PD neuron in each cell in the triangulation. (b) Distribution of absolute triadic differences over the entire dataset (gray), compared to triadic differences between shuffled triads (green) and compared to triadic differences between the first two principal components. Triadic differences are significantly smaller than in the shuffled data and in the principal components (*p < .*00fi, K-S test). (c-d) Same as in (a-b), for the duty cycle of the PD neuron.

**Figure 2–Figure supplement 3.**
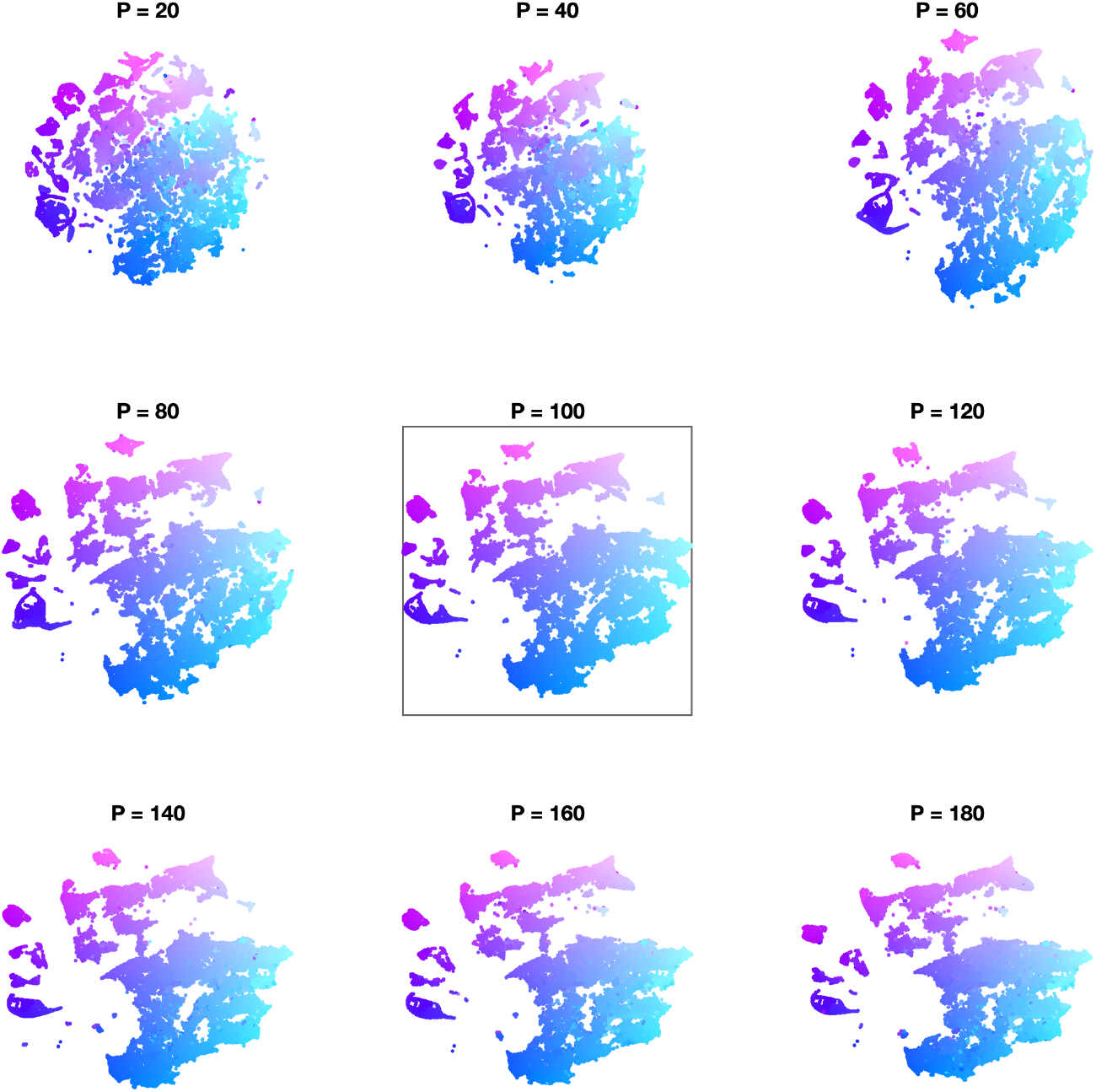
Effect of varying perplexity in t-SNE embedding. In each panel, data are embedded using t-SNE using the indicated perplexity parameter. Initialization is the same across all perplexities (Methods and Materials). In every panel, points are colored by their location in the embedding with *P* = l00 (black box).

**Figure 3–Figure supplement 1.**
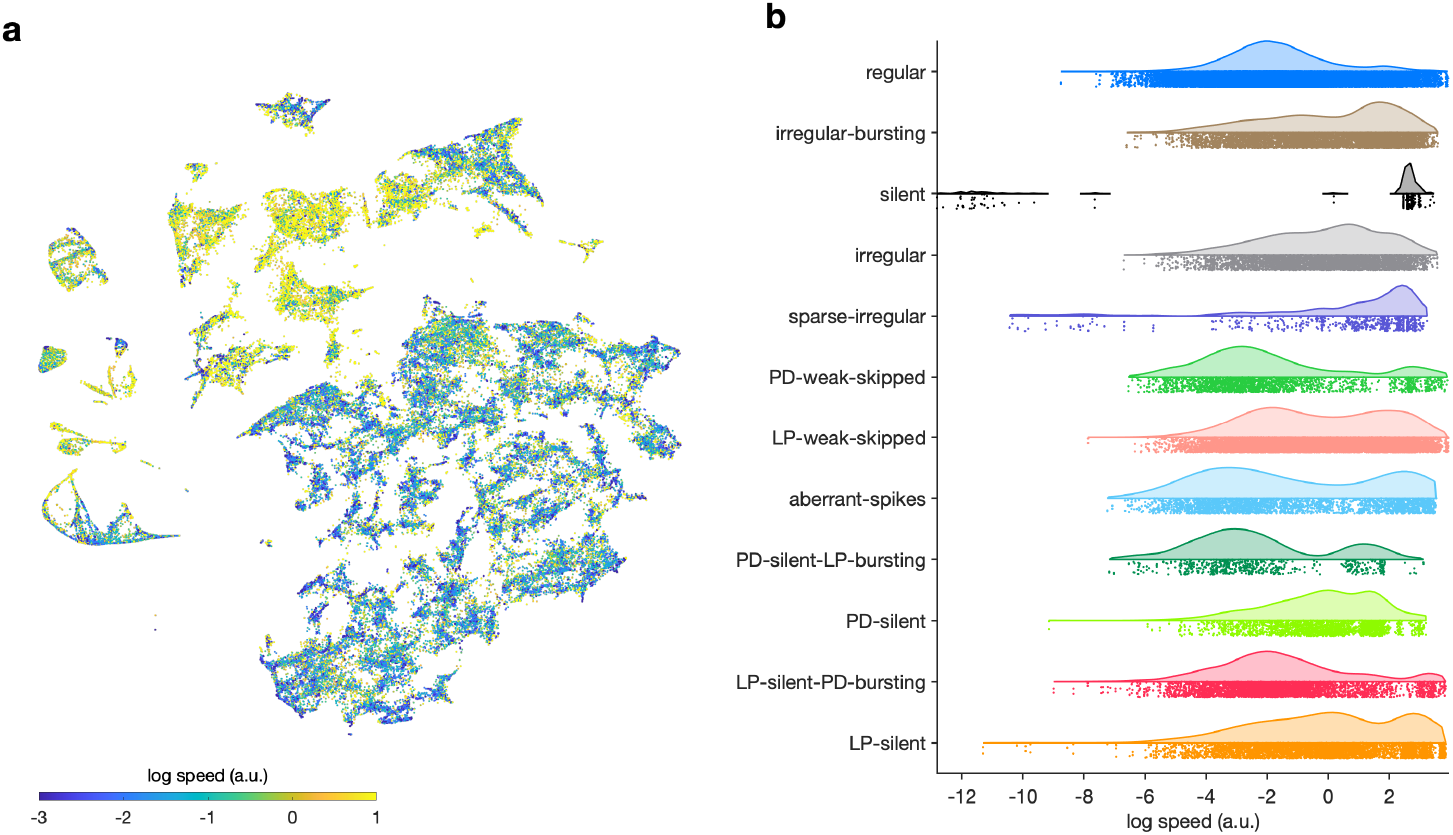
Speed of trajectories through map. (a) Map colored by speed of trajectories through map at that point. Cooler colors indicate that preparations move through that region of space more slowly and warmer colors suggest that preparations are more likely to be far from that location in the next time step. (b) Speed grouped by manually identified labels.

**Figure 3–Figure supplement 2.**
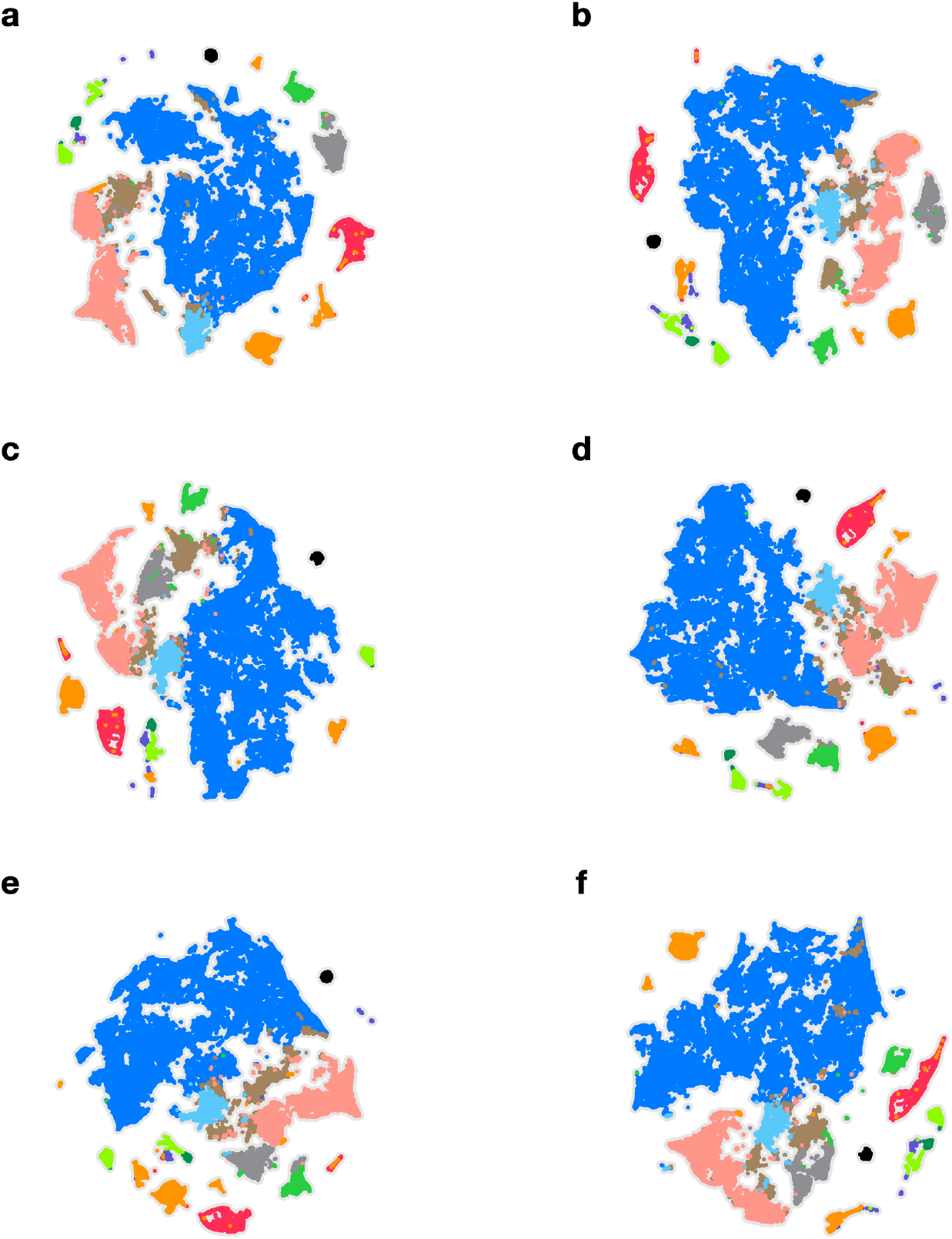
Embeddings with different initializations. In each panel, the embedding is performed with a different initialization. (a-d) Random initializations. (e) Initializations based on minimum ISIs in PD and LP. (f) Initializations based on mean ISIs in PD and LP. In every panel, points are colored identically.

**Figure 4–Figure supplement 1.**
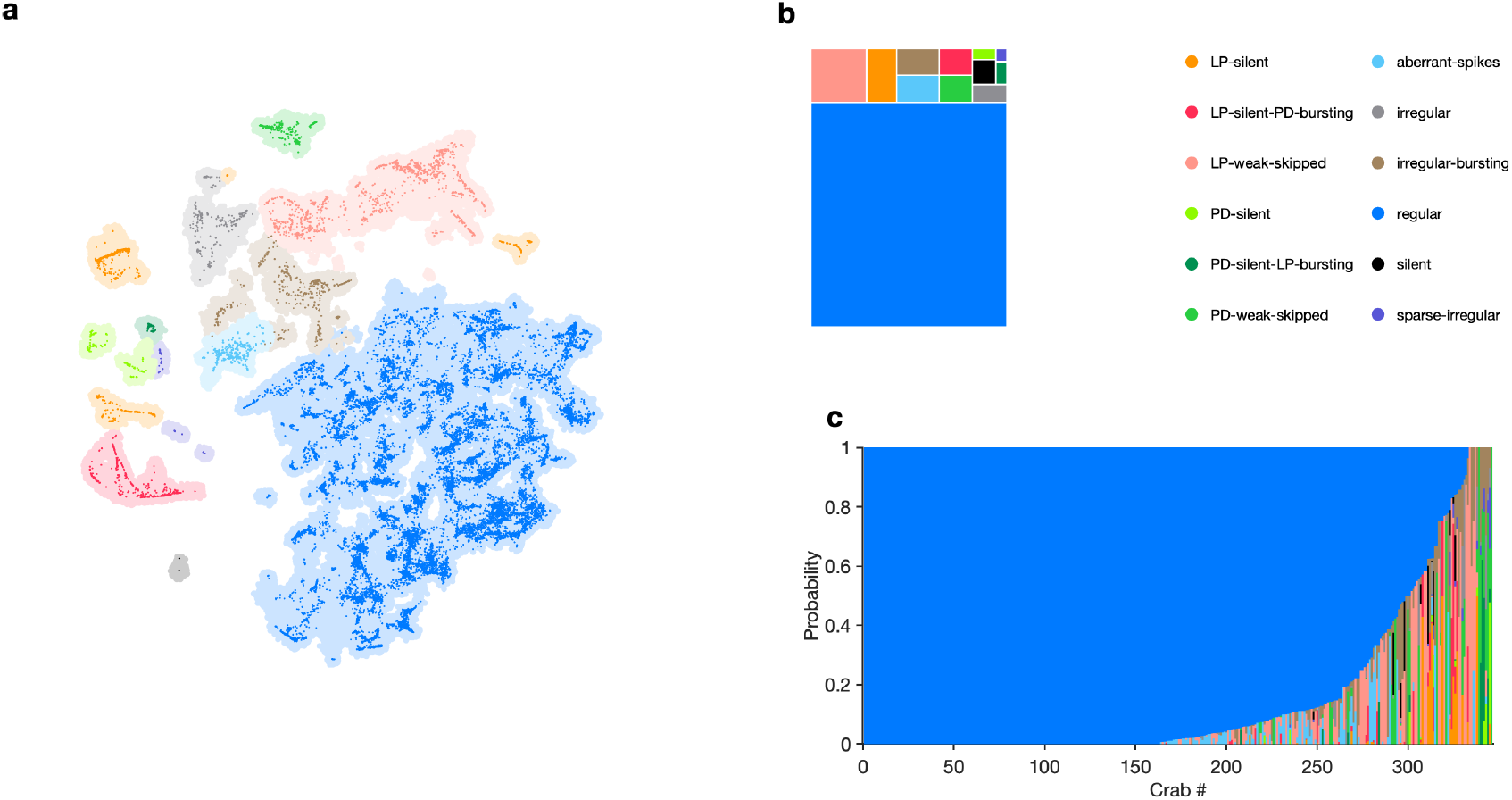
State distribution under baseline conditions. (a) Map showing occupancy of baseline data. Shading indicates all data. Bright colored points are data from baseline conditions. (b) Treemap showing state probabilities under baseline conditions. (c) Preparation-by-preparation variation in state distribution under baseline conditions. *n* = 22807 data snippets from *N* = 346 individual preparations.

**Figure 4–Figure supplement 2.**
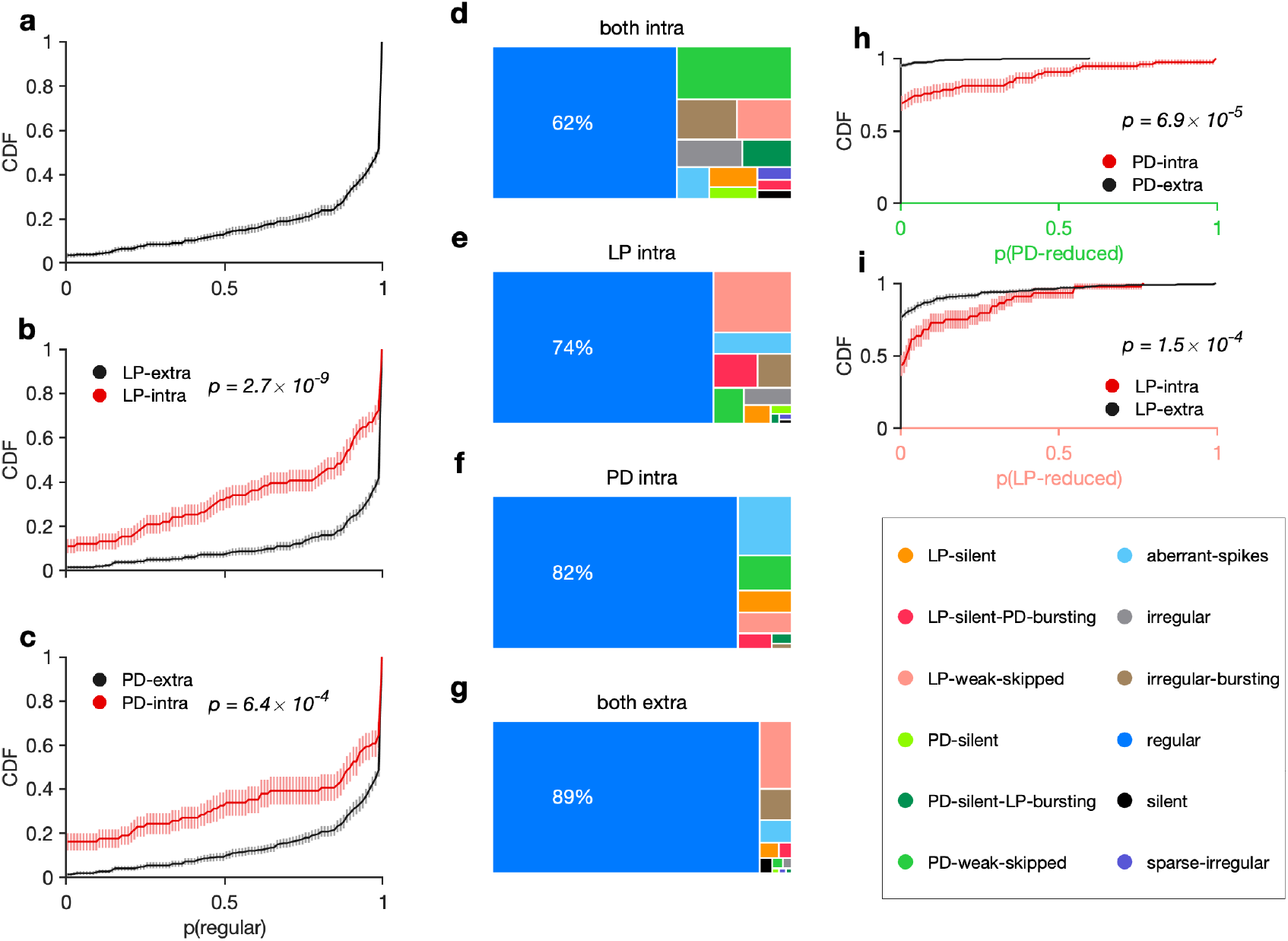
Recording condition alters regular state probability. (a) Cumulative probability of the probability of observing a regular state, per animal. Cumulative probability of probability of regular state for preparations with LP (b) and PD (c) recorded extracellularly (black) vs. LP recorded intracellularly (red). (d-g) State probabilities in different recording conditions. (h) Probability of observing states in which PD activity is reduced (PD-silent, PD-weak-skipped) in preparations in which PD is recorded intracellularly (red) vs. preparations in which PD is recorded extracellularly (black). (i) Same as (h), but for LP. In all panels, thick lines show CDFs and shading indicates confidence intervals estimated by bootstrap. *n* = 22807, *N* = 346. *p*-values in each panel are from two-sample Kolmogorov-Smirnoff tests.

**Figure 4–Figure supplement 3.**
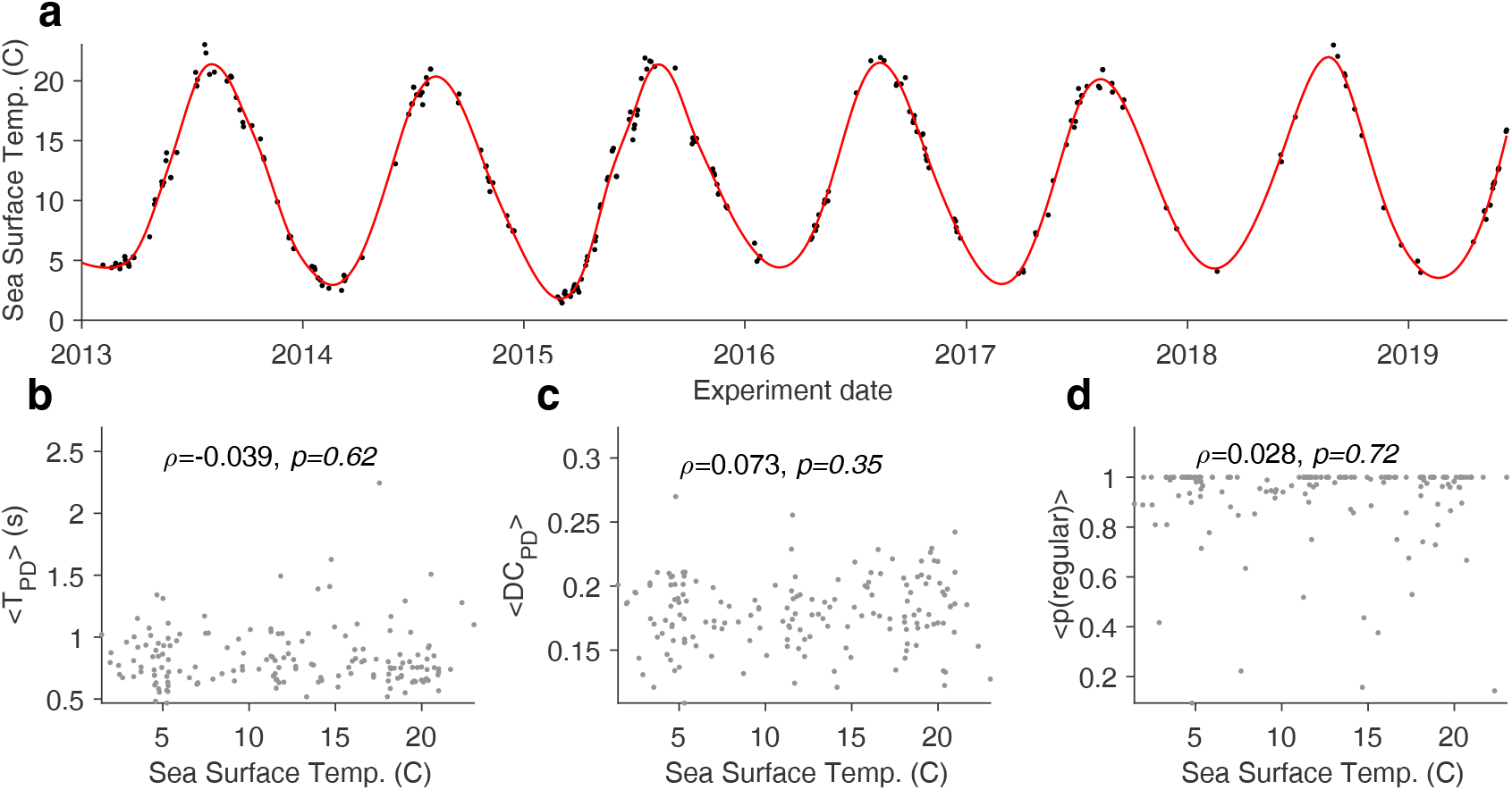
Effect of sea surface temperature on baseline circuit dynamics. (a) Sea surface temperature at the Boston Harbor vs. experimental date. Red line is a smoothing fit. Mean burst period of PD neuron (b), mean duty cycle of PD (c), and probability of observing the regular state (d) vs. sea surface temperature. In all panels, each dot corresponds to a single preparation. *N* = 312 preparations. *p* is the Spearman correlation coefficient and *p*-value is from the Spearman rank correlation test.

**Figure 5–Figure supplement 1.**
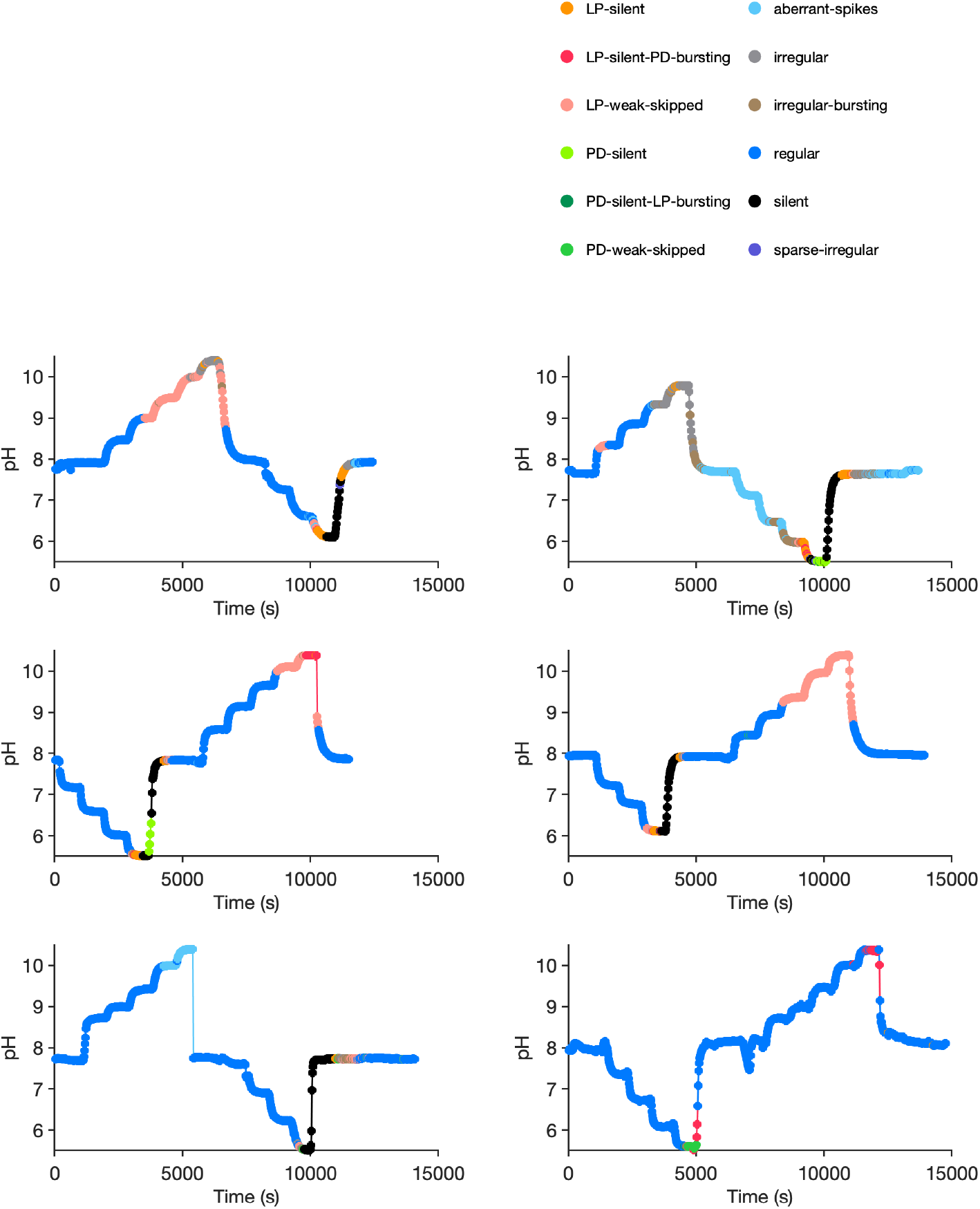
Preparation-by-preparation response to pH perturbations. Each panel shows the response of a single preparation to pH perturbations. States are indicated in colors. Each preparation was stepped through various pH levels before returning to baseline pH. Note silent states (black) during acidic pH.

**Figure 7–Figure supplement 1.**
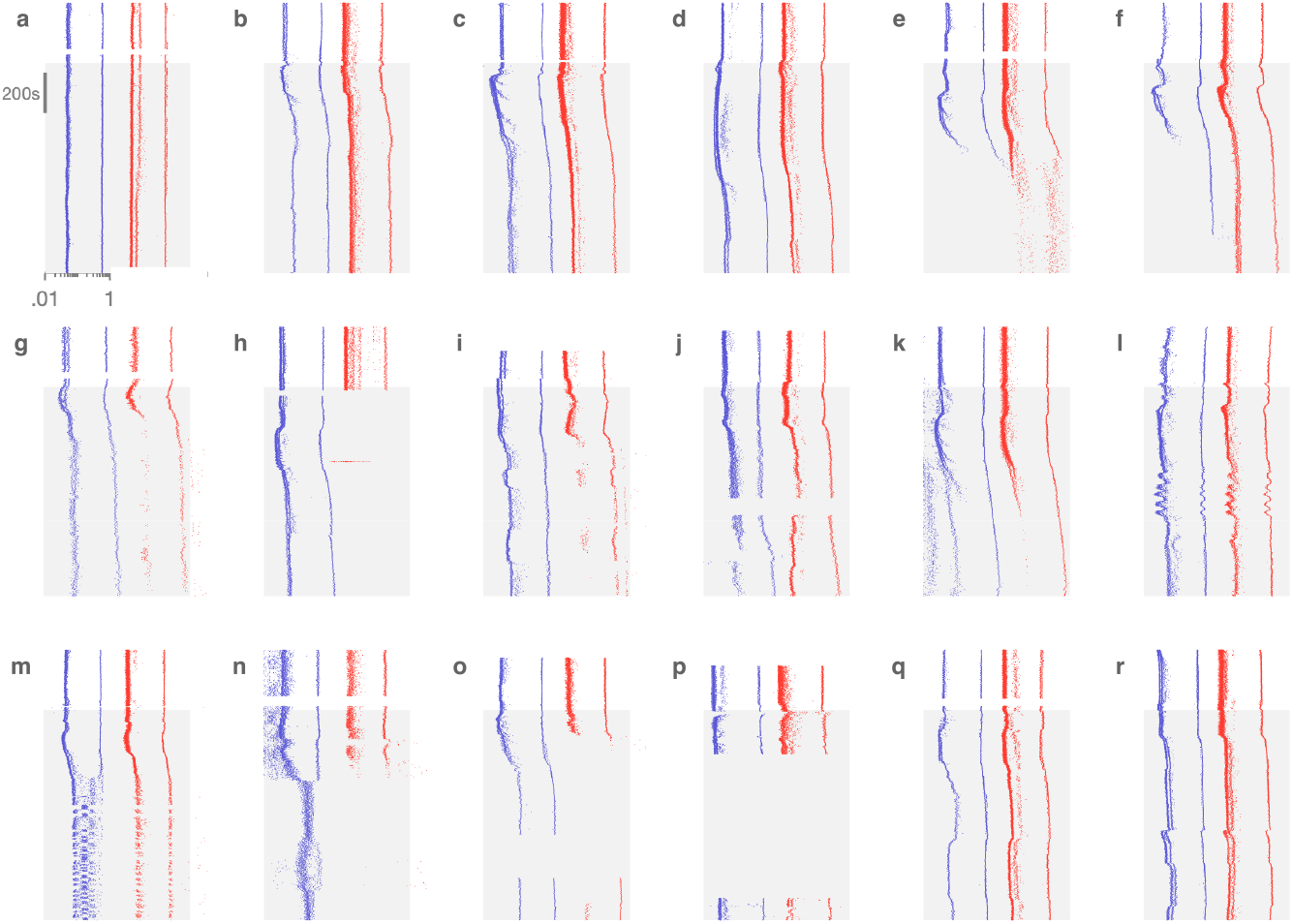
Decentralization evokes variable dynamics. In each panel, inter-spike intervals (ISIs) of PD (blue) and LP are shown before and after (shaded) decentralization. The diversity of circuit responses to decentralization include minimal change (a), transient perturbation followed by recovery (b-d), silence in one or two neurons (e-h), slow oscillatory responses (l) and a switch from bursting to spiking (m,n). Each panel corresponds to a different preparation.

**Figure 7–Figure supplement 2.**
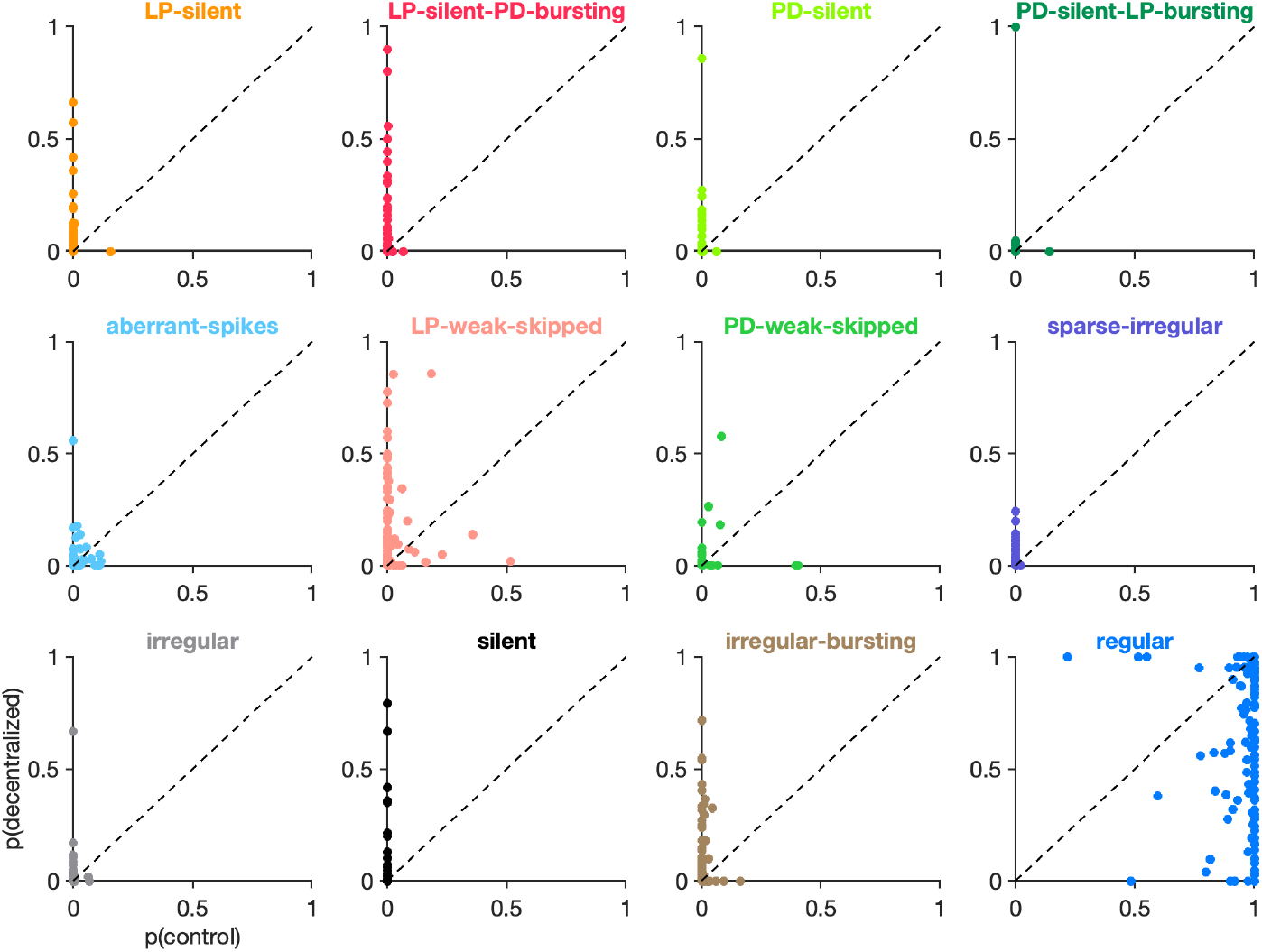
Effects of decentralization on state probabilities. Each panel shows the probability of observing a given state before (x-axis) and after (y-axis) decentralization. Each dot is a single preparation. Probabilities computed on an animal-by-animal basis from *n* = 16940 points from *N* = 140 preparations.

**Figure 7–Figure supplement 3.**
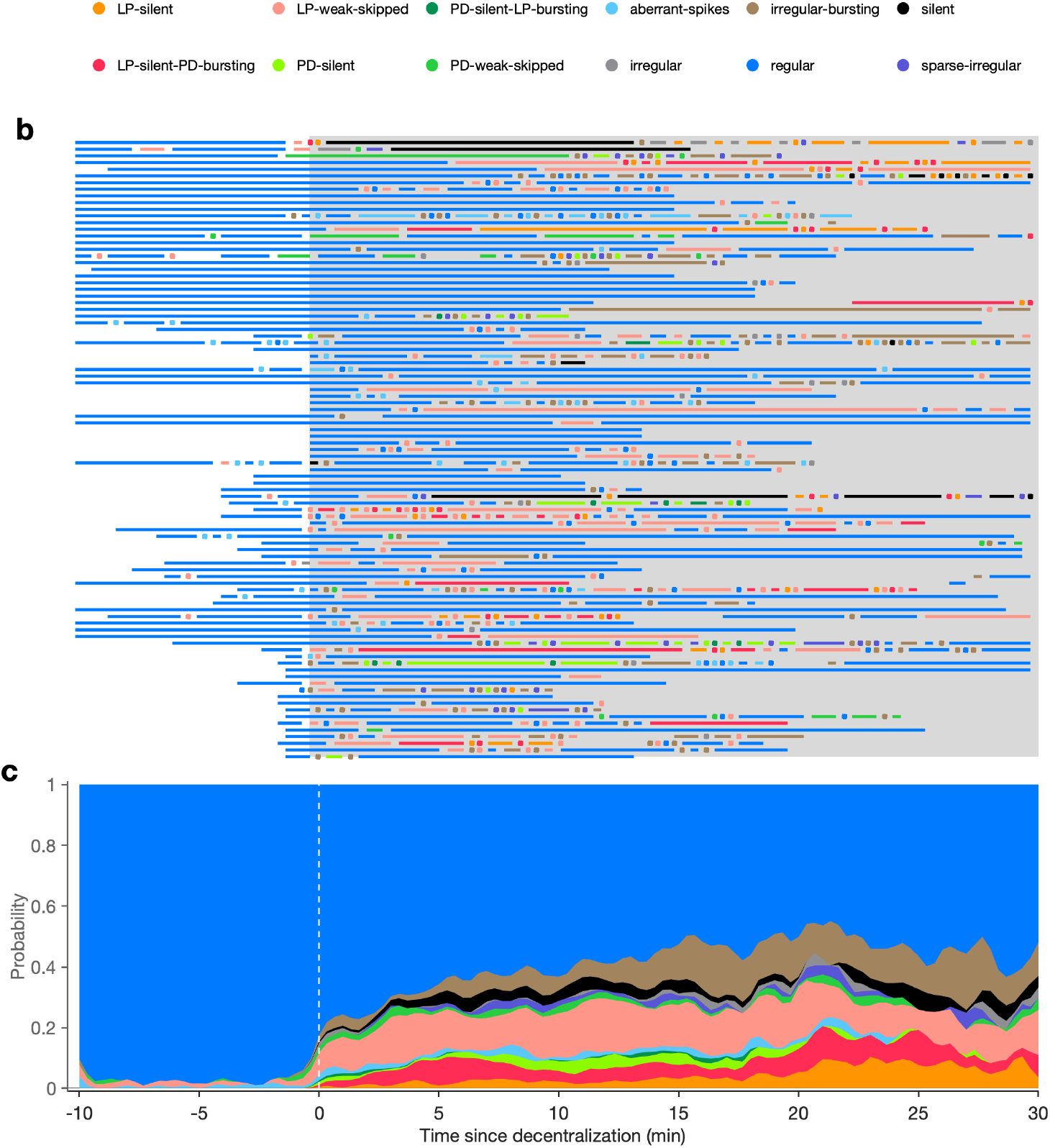
Time course of effects of decentralization. (a) Each line shows the states exhibited by one circuit before and after (gray shaded region) decentralization. Dots indicate states that were maintained only for one time bin (20s). (b) Stacked bars show probabilities of displaying state vs. time. *N* = 93 animals.

**Figure 7–Figure supplement 4.**
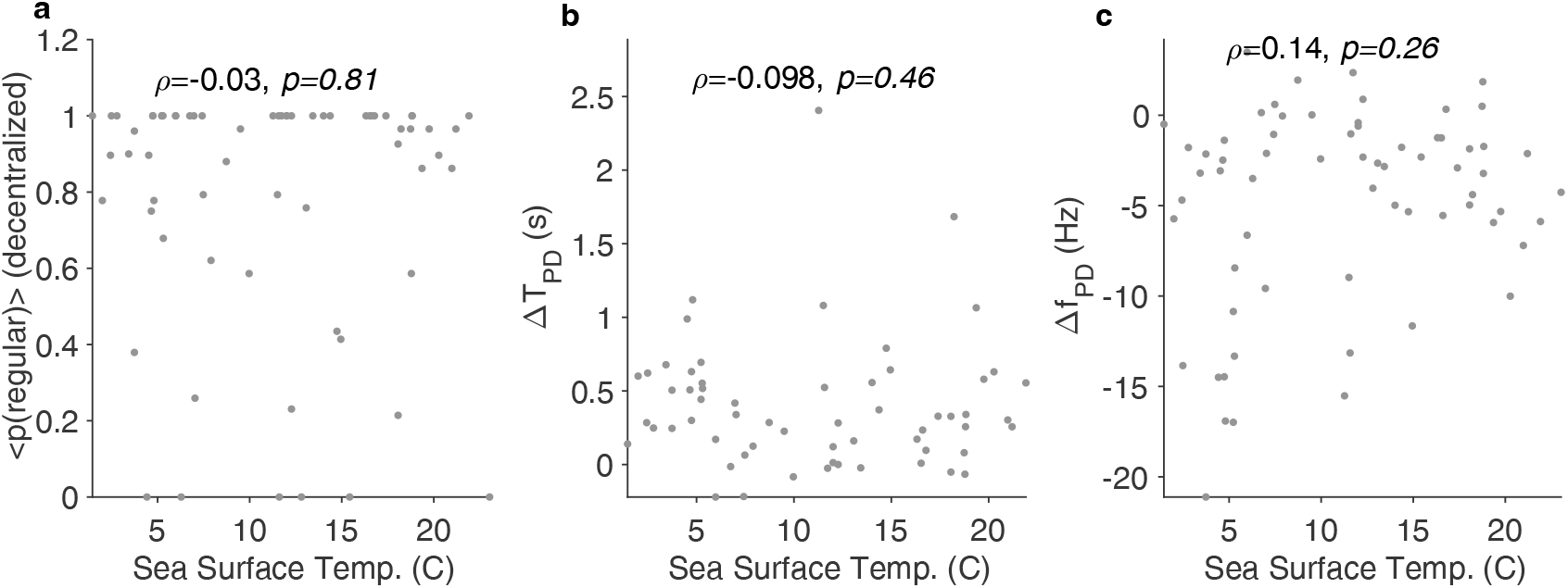
Effects of decentralization do not correlate with seasonal effects. Probability of observing the regular state during decentralization (a), change in time period of PD neuron (b), and change in firing rate of PD (c) vs. sea surface temperature on day experiment was carried out. *p* is the Spearman correlation coefficient, and *p*-values are computed using the Spearman rank correlation test.

**Figure 8–Figure supplement 1.**
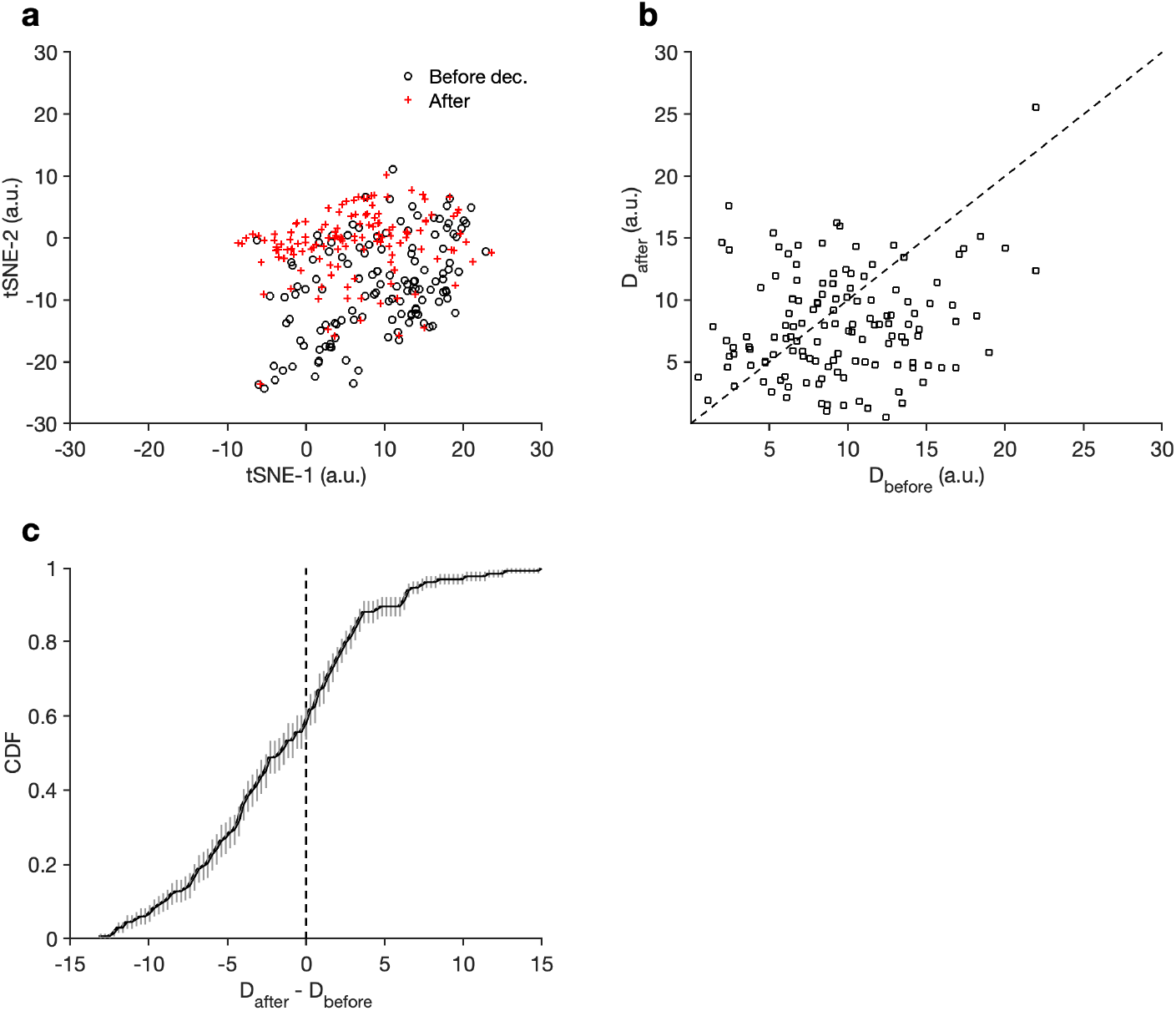
Effects of decentralization on regular rhythms. (a) Mean locations of data in the regular cluster before and after decentralization. Each preparation is represented by a pair of points, one circle and one cross. (b) Dispersion (Methods and Materials) of data before and after decentralization. Each preparation is a single point. Note that the data appear to be skewed to the right, indicating larger dispersion before decentralization. (c) Distribution of differences in dispersion. The distribution of differences is not significantly skewed from a Gaussian (*p* = .66, Anderson-Darling test), and dispersion in decentralized preparations is significantly lower than in baseline (*p* = .0016*, t* = 3.246, paired *t*-test). *n* = 13758 points from *N* = 140 preparations.

**Figure 9–Figure supplement 1.**
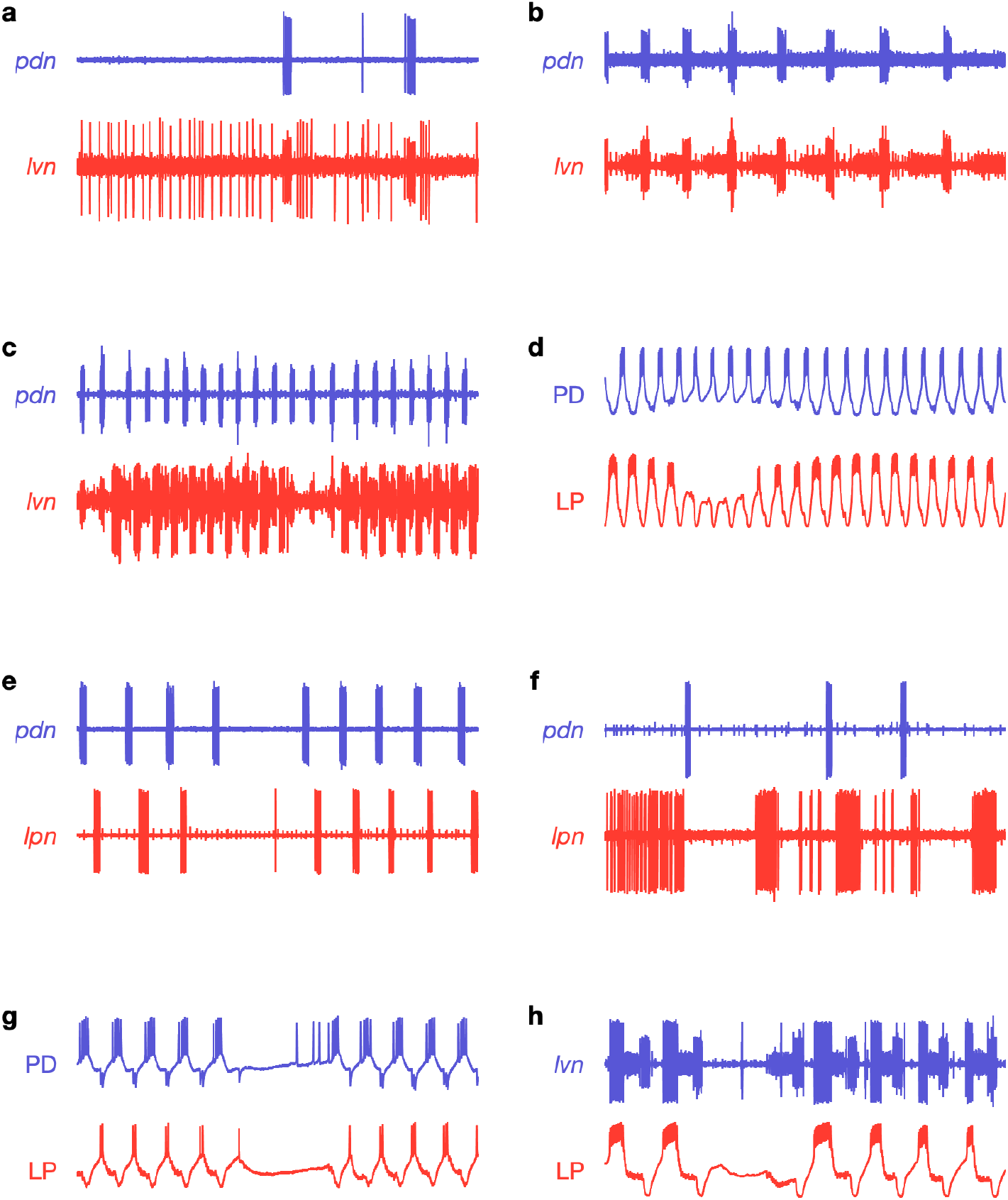
Non-regular activity patterns in proctolin. Each panel shows a 20s snippet of raw recordings showing spikes from LP (red) and PD (blue). Each panel is from a different animal. Each row is from a different experimenter. (a) Irregular bursting, note prolonged spiking of LP on *lvn*. (b) LP completely silent, missing from *lvn*. (c) Intermittent LP interruptions, note breaks in *lvn*. (d) Interruption in LP bursting. (e) Interruption in PD and LP bursting. (f) Irregular bursting of both PD and LP. (g-h) Interruption of both PD and LP. Traces labelled PD or LP are intracellular recordings.

**Figure 9–Figure supplement 2.**
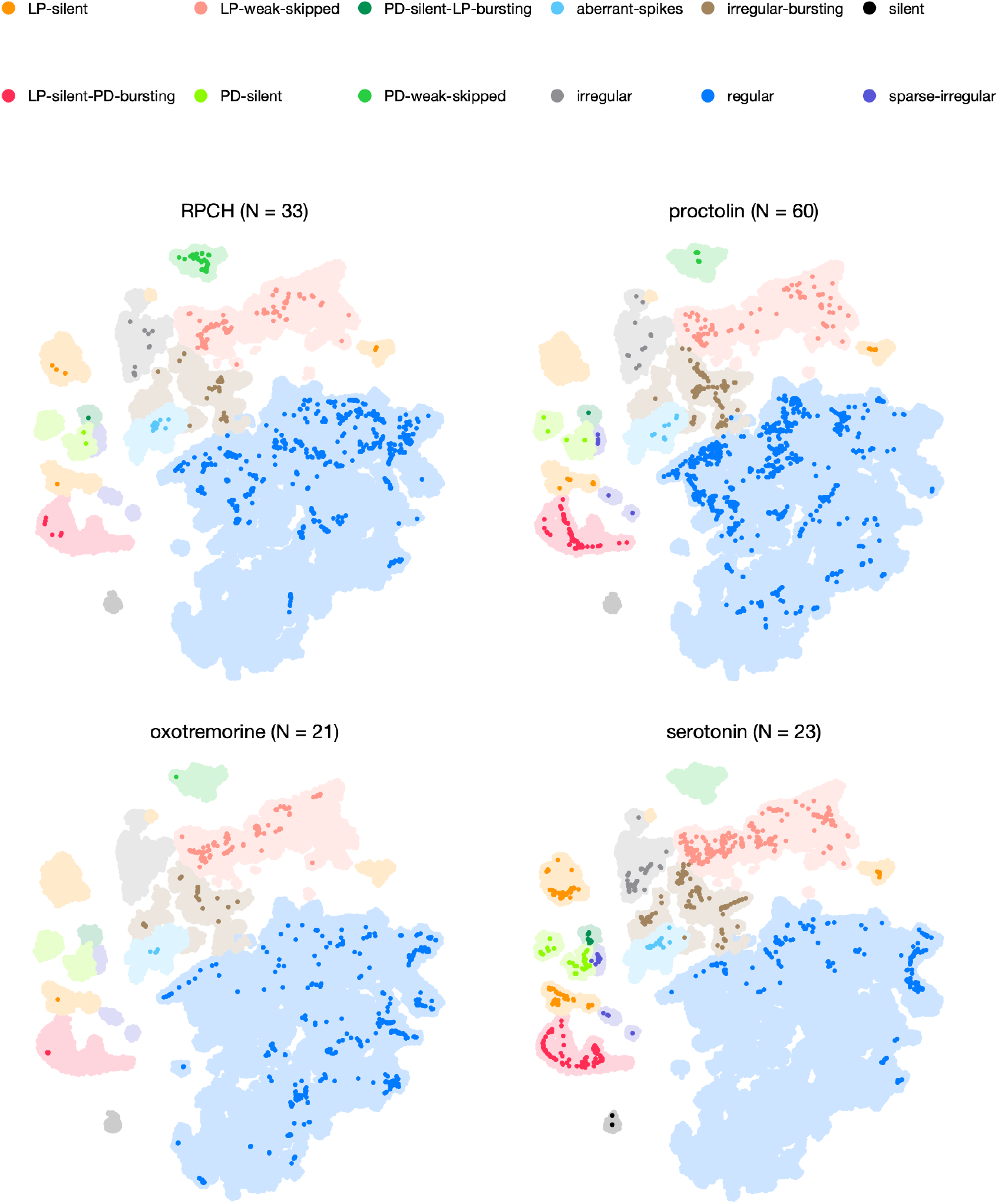
Neuromodulators affect map occupancy. In each panel, all the data are shown in light shading. Bright colors indicate distribution of data during bath application of that neuromodulator. The number of animals in each panel is indicated in the title.

## Notes

### Competing Interest Statement

The authors have declared no competing interest.

## References

Allen M, Poggiali D, Whitaker K, Marshall TR, Kievit RA. Raincloud plots: a multi-platform tool for robust data visualization. Wellcome open research. 2019; 4.

Alonso LM, Marder E. Visualization of currents in neural models with similar behavior and different conductance densities. eLife. 2019; 8:e42722.

Alonso LM, Marder E. Temperature compensation in a small rhythmic circuit. eLife. 2020; 9:e55470.

Bal T, Nagy F, Moulins M. Muscarinic modulation of a pattern-generating network: control of neuronal properties. Journal of Neuroscience. 1994; 14(5):3019–3035.

Baldi P. Autoencoders, unsupervised learning, and deep architectures. In: Proceedings of ICML workshop on unsupervised and transfer learning JMLR Workshop and Conference Proceedings; 2012. p. 37–49.

Beltz B, Eisen JS, Flamm R, Harris-Warrick R, Hooper SL, Marder E. Serotonergic innervation and modulation of the stomatogastric ganglion of three decapod crustaceans (*Panulirus interruptus*, Homarus americanus and Cancer irroratus). Journal of experimental biology. 1984; 109(1):35–54.

Berman GJ, Choi DM, Bialek WS, Shaevitz JW. Mapping the stereotyped behaviour of freely moving fruit flies. Journal of the Royal Society Interface. 2014 Oct; 11(99):20140672–20140672.

Böhm JN, Berens P, Kobak D. A Unifying Perspective on Neighbor Embeddings along the Attraction-Repulsion Spectrum. arXiv preprint arXiv:200708902. 2020; .

Börner K, Chen C, Boyack KW. Visualizing knowledge domains. Annual review of information science and technology. 2003; 37(1):179–255.

Brunel N, Wang XJ. What determines the frequency of fast network oscillations with irregular neural discharges? I. Synaptic dynamics and excitation-inhibition balance. Journal of Neurophysiology. 2003; 90(1):415–430.

Bucher D, Prinz AA, Marder E. Animal-to-animal variability in motor pattern production in adults and during growth. Journal of Neuroscience. 2005 Feb; 25(7):1611–1619.

Caplan JS, Williams AH, Marder E. Many Parameter Sets in a Multicompartment Model Oscillator Are Robust to Temperature Perturbations. Journal of Neuroscience. 2014 Apr; 34(14):4963–4975.

Chen W, Zhao Y, Chen X, Yang Z, Xu X, Bi Y, Chen V, Li J, Choi H, Ernest B, Tran B, Mehta M, Kumar P, Farmer A, Mir A, Mehra UA, Li JL, Moos M, Xiao W, Wang C. A multicenter study benchmarking single-cell RNA sequencing technologies using reference samples. Nature Biotechnology. 2020 Dec; p. 1–29.

Christen M, Kohn A, Ott T, Stoop R. Measuring spike pattern reliability with the Lempel–Ziv-distance. Journal of Neuroscience Methods. 2006; 156(1-2):342–350.

Clark DG, Livezey JA, Bouchard KE. Unsupervised Discovery of Temporal Structure in Noisy Data with Dynamical Components Analysis. arXiv. 2019 May; .

Clark MC. Arthropod 5-HT2 Receptors: A Neurohormonal Receptor in Decapod Crustaceans That Displays Agonist Independent Activity Resulting from an Evolutionary Alteration to the DRY Motif. Journal of Neuroscience. 2004 Mar; 24(13):3421–3435.

Clemens S, Massabuau JC, Meyrand P, Simmers J. A modulatory role for oxygen in shaping rhythmic motor output patterns of neuronal networks. Respir Physiol. 2001 Nov; 128(3):299–315.

Corver A, Wilkerson N, Miller J, Gordus AG. Distinct movement patterns generate stages of spider web-building. bioRxiv. 2021; .

Cox MA, Cox TF. Multidimensional scaling. In: Handbook of data visualization Springer; 2008.p. 315–347.

Cunningham JP, Yu BM. Dimensionality reduction for large-scale neural recordings. Nature Neuroscience. 2014 Aug; 17(11):1500–1509.

Cymbalyuk GS, Gaudry Q, Masino MA, Calabrese RL. Bursting in leech heart interneurons: cell-autonomous and network-based mechanisms. Journal of Neuroscience. 2002 Dec; 22(24):10580–10592.

Daur N, Nadim F, Bucher D. The complexity of small circuits: the stomatogastric nervous system. Current Opinion in Neurobiology. 2016; 41:1–7.

Dickinson PS, Hauptman J, Hetling J, Mahadevan A. RPCH modulation of a multi-oscillator network: effects on the pyloric network of the spiny lobster. Journal of Neurophysiology. 2001; 85(4):1424–1435.

Dimitriadis G, Neto JP, Kampff AR. t-SNE Visualization of Large-Scale Neural Recordings. Neural Computation. 2018 Jun; 30(7):1750–1774.

Eisen JS, Marder E. A mechanism for production of phase shifts in a pattern generator. Journal of neurophysiology. 1984; 51(6):1375–1393.

Eisen JS, Marder E. Mechanisms underlying pattern generation in lobster stomatogastric ganglion as determined by selective inactivation of identified neurons. III. Synaptic connections of electrically coupled pyloric neurons. Journal of Neurophysiology. 1982; 48(6):1392–1415.

Epstein I, Marder E. Multiple modes of a conditional neural oscillator. Biological cybernetics. 1990; 63(1):25–34.

Fortuin V, Hüser M, Locatello F, Strathmann H, Rätsch G. SOM-VAE: Interpretable Discrete Representation Learning on Time Series. arXiv. 2018 Jun; .

Franci A, O’Leary T, Golowasch J. Positive dynamical networks in neuronal regulation: how tunable variability coexists with robustness. IEEE Control Systems Letters. 2020; .

Frankel NW, Pontius W, Dufour YS, Long J, Hernandez-Nunez L, Emonet T. Adaptability of non-genetic diversity in bacterial chemotaxis. eLife. 2014; 3:e03526.

Garcia VJ, Daur N, Temporal S, Schulz DJ, Bucher D. Neuropeptide Receptor Transcript Expression Levels and Magnitude of Ionic Current Responses Show Cell Type-Specific Differences in a Small Motor Circuit. Journal of Neuroscience. 2015 Apr; 35(17):6786–6800.

Golowasch J, Marder E. Proctolin activates an inward current whose voltage dependence is modified by extracellular *Ca*^2+^. Journal of Neuroscience. 1992 Mar; 12(3):810–817.

Golowasch J, Casey M, Abbott L, Marder E. Network Stability from Activity-Dependent Regulation of Neuronal Conductances. Neural Computation. 1999 Jul; 11(5):1079–1096.

Golowasch J, Goldman MS, Abbott L, Marder E. Failure of Averaging in the Construction of a Conductance-Based Neuron Model. Journal of Neurophysiology. 2002 Feb; 87(2):1129–1131.

Gonçalves PJ, Lueckmann JM, Deistler M, Nonnenmacher M, Öcal K, Bassetto G, Chintaluri C, Podlaski WF, Haddad SA, Vogels TP, et al. Training deep neural density estimators to identify mechanistic models of neural dynamics. eLife. 2020; 9:e56261.

Gorur-Shandilya S, Marder E, O’Leary T. Activity-dependent compensation of cell size is vulnerable to targeted deletion of ion channels. Scientific Reports. 2020; 10.

Gutierrez GJ, Grashow RG. *Cancer borealis* stomatogastric nervous system dissection. Journal of visualized experiments: JoVE. 2009; (25).

Gutierrez GJ, O’Leary T, Marder E. Multiple mechanisms switch an electrically coupled, synaptically inhibited neuron between competing rhythmic oscillators. Neuron. 2013; 77(5):845–858.

Haddad SA, Marder E. Circuit Robustness to Temperature Perturbation Is Altered by Neuromodulators. Neuron. 2018 Nov; 100(3):609–623.

Haley JA, Hampton D, Marder E. Two central pattern generators from the crab, *Cancer borealis*, respond robustly and differentially to extreme extracellular pH. eLife. 2018 Dec; 7:e41877.

Hamood AW, Haddad SA, Otopalik AG, Rosenbaum P, Marder E. Quantitative Reevaluation of the Effects of Short- and Long-Term Removal of Descending Modulatory Inputs on the Pyloric Rhythm of the Crab, *Cancer borealis*. eNeuro. 2015 Jan; 2(1):1–13.

Hamood AW, Marder E. Animal-to-Animal Variability in Neuromodulation and Circuit Function. Cold Spring Harb Symp Quant Biol. 2015 Jun; 79:21–28.

Harris-Warrick RM, Flamm RE. Multiple mechanisms of bursting in a conditional bursting neuron. Journal of Neuroscience. 1987; 7(7):2113–2128.

Harris-Warrick RM, Marder E. Modulation of neural networks for behavior. Annual review of neuroscience. 1991; 14(1):39–57.

Hartline DK, Maynard DM. Motor patterns in the stomatogastric ganglion of the lobster *Panulirus argus*. Journal of Experimental Biology. 1975; 62(2):405–420.

He LS, Rue MC, Morozova EO, Powell DJ, James EJ, Kar M, Marder E. Rapid adaptation to elevated extracellular potassium in the pyloric circuit of the crab, *Cancer borealis*. Journal of Neurophysiology. 2020; 123(5):2075–2089.

Hooper SL, Marder E. Modulation of a central pattern generator by two neuropeptides, proctolin and FMR-Famide. Brain Research. 1984 Jul; 305(1):186–191.

Hooper SL, Marder E. Modulation of the lobster pyloric rhythm by the peptide proctolin. Journal of Neuroscience. 1987; 7(7):2097–2112.

Hooper SL, Thuma JB, Guschlbauer C, Schmidt J, Büschges A. Cell dialysis by sharp electrodes can cause nonphysiological changes in neuron properties. Journal of Neurophysiology. 2015; 114(2):1255–1271.

Kobak D, Berens P. The art of using t-SNE for single-cell transcriptomics. Nature Communications. 2019 Nov; p. 1–14.

Kobak D, Linderman GC. Initialization is critical for preserving global data structure in both t-SNE and UMAP. Nature biotechnology. 2021; 39(2):156–157.

Kollmorgen S, Hahnloser RH, Mante V. Nearest neighbours reveal fast and slow components of motor learning. Nature. 2020; 577(7791):526–530.

Kushinsky D, Morozova EO, Marder E. In vivo effects of temperature on the heart and pyloric rhythms in the crab *Cancer borealis*. Journal of Experimental Biology. 2019; 222(5).

Kyriazi P, Headley DB, Pare D. Multi-dimensional Coding by Basolateral Amygdala Neurons. Neuron. 2018 Sep; 99(6):1315–1328.e5.

Leelatian N, Sinnaeve J, Mistry AM, Barone SM, Brockman AA, Diggins KE, Greenplate AR, Weaver KD, Thompson RC, Chambless LB, Mobley BC, Ihrie RA, Irish JM. Unsupervised machine learning reveals risk stratifying glioblastoma tumor cells. eLife. 2020 Jun; 9:545.

Linderman GC, Rachh M, Hoskins JG, Steinerberger S, Kluger Y. Fast interpolation-based t-SNE for improved visualization of single-cell RNA-seq data. Nature methods. 2019; 16(3):243–245.

Linderman GC, Steinerberger S. Clustering with t-SNE, provably. SIAM Journal on Mathematics of Data Science. 2019; 1(2):313–332.

Van der Maaten L, Hinton G. Visualizing data using t-SNE. Journal of machine learning research. 2008; 9(11).

Mackevicius EL, Bahle AH, Williams AH, Gu S, Denisenko NI, Goldman MS, Fee MS. Unsupervised discovery of temporal sequences in high-dimensional datasets, with applications to neuroscience. eLife. 2019; 8:e38471.

Macosko EZ, Basu A, Satija R, Nemesh J, Shekhar K, Goldman M, Tirosh I, Bialas AR, Kamitaki N, Martersteck EM, Trombetta JJ, Weitz DA, Sanes JR, Shalek AK, Regev A, McCarroll SA. Highly Parallel Genome-wide Expression Pro1ling of Individual Cells Using Nanoliter Droplets. Cell. 2015 May; 161(5):1202–1214.

Madiraju NS, Sadat SM, Fisher D, Karimabadi H. Deep Temporal Clustering : Fully Unsupervised Learning of Time-Domain Features. arXiv. 2018 Feb; .

Marbán E. Cardiac channelopathies. Nature. 2002 Jan; 415(6868):213–218.

Marder E. Neuromodulation of neuronal circuits: back to the future. Neuron. 2012; 76(1):1–11.

Marder E, Bucher D. Understanding circuit dynamics using the stomatogastric nervous system of lobsters and crabs. Annu Rev Physiol. 2007; 69:291–316.

Marder E, Hooper SL. Neurotransmitter modulation of the stomatogastric ganglion of decapod crustaceans. In: Model neural networks and behavior Springer; 1985. p. 319–337.

Marder E, Hooper SL, Siwicki KK. Modulatory action and distribution of the neuropeptide proctolin in the crustacean stomatogastric nervous system. Journal of Comparative Neurology. 1986; 243(4):454–467.

Marder E, Weimann JM. Modulatory control of multiple task processing in the stomatogastric nervous system. In: Neurobiology of motor programme selection Elsevier; 1992.p. 3–19.

Mariño J, Schummers J, Lyon DC, Schwabe L, Beck O, Wiesing P, Obermayer K, Sur M. Invariant computations in local cortical networks with balanced excitation and inhibition. Nature neuroscience. 2005; 8(2):194–201.

Miller JP, Selverston AI. Mechanisms underlying pattern generation in lobster stomatogastric ganglion as determined by selective inactivation of identified neurons. IV. Network properties of pyloric system. Journal of neurophysiology. 1982; 48(6):1416–1432.

Mizrahi A, Dickinson PS, Kloppenburg P, Fénelon V, Baro DJ, Harris-Warrick RM, Meyrand P, Simmers J. Longterm maintenance of channel distribution in a central pattern generator neuron by neuromodulatory inputs revealed by decentralization in organ culture. Journal of Neuroscience. 2001; 21(18):7331–7339.

Moor M, Horn M, Rieck B, Borgwardt K. Topological Autoencoders. arXiv. 2019 Jun; .

Nguyen LH, Holmes S. Ten quick tips for effective dimensionality reduction. PLoS Computational Biology. 2019 Jun; 15(6):e1006907–19.

Nusbaum MP, Marder E. A Neuronal Role for a Crustacean Red Pigment Concentrating Hormone-like Peptide: Neuromodulation of the Pyloric Rhythm in the Crab, *Cancer Borealis*. Journal of Experimental Biology. 1988 Sep; 135:1–17.

Nusbaum MP, Marder E. A modulatory proctolin-containing neuron (MPN). I. Identification and characterization. Journal of Neuroscience. 1989; 9(5):1591–1599.

O’Leary T, Williams AH, Franci A, Marder E. Cell Types, Network Homeostasis, and Pathological Compensation from a Biologically Plausible Ion Channel Expression Model. Neuron. 2014 May; 82(4):809–821.

Pang R, Lansdell BJ, Fairhall AL. Dimensionality reduction in neuroscience. Current Biology. 2016 Jul; 26(14):R656–60.

Peacock JA. Two-dimensional goodness-of-fit testing in astronomy. Monthly Notices of the Royal Astronomical Society. 1983; 202(3):615–627.

Powell D, Haddad SA, Gorur-Shandilya S, Marder E. Coupling between fast and slow oscillator circuits in *Cancer borealis* is temperature-compensated. eLife. 2021; 10:e60454.

Prinz AA, Billimoria CP, Marder E. Alternative to Hand-Tuning Conductance-Based Models: Construction and Analysis of Databases of Model Neurons. Journal of Neurophysiology. 2003 Dec; 90(6):3998–4015.

Prinz AA, Bucher D, Marder E. Similar network activity from disparate circuit parameters. Nature Neuroscience. 2004 Nov; 7(12):1345–1352.

Qadri SA, Camacho J, Wang H, Taylor JR, Grosell M, Worden MK. Temperature and acid–base balance in the American lobster *Homarus americanus*. Journal of Experimental Biology. 2007; 210(7):1245–1254.

Ratliff J, Franci A, Marder E, O’Leary T. Neuronal oscillator robustness to multiple global perturbations. Biophysical Journal. 2021; .

Rinberg A, Taylor AL, Marder E. The effects of temperature on the stability of a neuronal oscillator. PLoS Computational Biology. 2013; 9(1):e1002857.

Rosenbaum P, Marder E. Graded Transmission without Action Potentials Sustains Rhythmic Activity in Some But Not All Modulators That Activate the Same Current. Journal of Neuroscience. 2018 Oct; 38(42):8976–8988.

Rossum Mv. A novel spike distance. Neural computation. 2001; 13(4):751–763.

Rumelhart DE, Hinton GE, Williams RJ. Learning internal representations by error propagation. California Univ San Diego La Jolla Inst for Cognitive Science; 1985.

Russell DF. Rhythmic excitatory inputs to the lobster stomatogastric ganglion. Brain research. 1976; 101(3):582–588.

Schreiber S, Fellous JM, Whitmer D, Tiesinga P, Sejnowski TJ. A new correlation-based measure of spike timing reliability. Neurocomputing. 2003; 52:925–931.

Schulz DJ, Goaillard JM, Marder E. Variable channel expression in identified single and electrically coupled neurons in different animals. Nature neuroscience. 2006; 9(3):356–362.

Schulz DJ, Goaillard JM, Marder EE. Quantitative expression profiling of identified neurons reveals cell-specific constraints on highly variable levels of gene expression. Proceedings of the National Academy of Sciences. 2007; 104(32):13187–13191.

Settles B. Active learning literature survey. Doctoral Dissertation, University of Wisconsin-Madison. 2009; .

Shneiderman B, Wattenberg M. Ordered treemap layouts. In: IEEE Symposium on Information Visualization, 2001. INFOVIS 2001. IEEE; 2001. p. 73–78.

Spitzer N, Cymbalyuk G, Zhang H, Edwards DH, Baro DJ. Serotonin Transduction Cascades Mediate Variable Changes in Pyloric Network Cycle Frequency in Response to the Same Modulatory Challenge. Journal of Neurophysiology. 2008 Jun; 99(6):2844–2863.

Staley K. Molecular mechanisms of epilepsy. Nature Neuroscience. 2015 Feb; 18(3):367–372.

Swensen AM, Marder E. Multiple peptides converge to activate the same voltage-dependent current in a central pattern-generating circuit. Journal of Neuroscience. 2000; 20(18):6752–6759.

Swensen AM, Marder E. Modulators with Convergent Cellular Actions Elicit Distinct Circuit Outputs. Journal of Neuroscience. 2001 Jun; 21(11):4050–4058.

Tang LS, Taylor AL, Rinberg A, Marder E. Robustness of a Rhythmic Circuit to Short- and Long-Term Temperature Changes. Journal of Neuroscience. 2012 Jul; 32(29):10075–10085.

Tang LS, Goeritz ML, Caplan JS, Taylor AL, Fişek M, Marder E. Precise Temperature Compensation of Phase in a Rhythmic Motor Pattern. PLoS Biology. 2010 Aug; 8(8):e1000469.

Thirumalai V, Marder E. Colocalized Neuropeptides Activate a Central Pattern Generator by Acting on Different Circuit Targets. Journal of Neuroscience. 2002 Mar; 22(5):1874–1882.

Thirumalai V, Prinz AA, Johnson CD, Marder E. Red Pigment Concentrating Hormone Strongly Enhances the Strength of the Feedback to the Pyloric Rhythm Oscillator But Has Little Effect on Pyloric Rhythm Period. Journal of Neurophysiology. 2006 Mar; 95(3):1762–1770.

Thoby-Brisson M, Simmers J. Neuromodulatory inputs maintain expression of a lobster motor pattern-generating network in a modulation-dependent state: evidence from long-term decentralization *in vitro*. Journal of Neuroscience. 1998; 18(6):2212–2225.

Timme M. Revealing network connectivity from response dynamics. Physical review letters. 2007; 98(22):224101.

Tobin AE, Cruz-Bermúdez ND, Marder E, Schulz DJ. Correlations in ion channel mRNA in rhythmically active neurons. PloS one. 2009; 4(8):e6742.

Turrigiano G, Abbott L, Marder E. Activity-dependent changes in the intrinsic properties of cultured neurons. Science. 1994 May; 264(5161):974–977.

Turrigiano G, LeMasson G, Marder E. Selective regulation of current densities underlies spontaneous changes in the activity of cultured neurons. Journal of Neuroscience. 1995 May; 15(5):3640–3652.

Turrigiano GG, Marder E. Modulation of identified stomatogastric ganglion neurons in primary cell culture. Journal of Neurophysiology. 1993 Jun; 69(6):1993–2002.

Victor JD, Purpura KP. Metric-space analysis of spike trains: theory, algorithms and application. Network: computation in neural systems. 1997; 8(2):127–164.

van Vreeswijk C, Sompolinsky H. Chaos in neuronal networks with balanced excitatory and inhibitory activity. Science. 1996 Dec; 274(5293):1724–1726.

Vyas S, Golub MD, Sussillo D, Shenoy KV. Computation Through Neural Population Dynamics. Annual Reviews of Neuroscience. 2020 Jul; 43(1):249–275.

Weimann JM, Marder E. Switching neurons are integral members of multiple oscillatory networks. Current Biology. 1994; 4(10):896–902.

Williams AH, Degleris A, Wang Y, Linderman SW. Point process models for sequence detection in high-dimensional neural spike trains. arXiv. 2020 Oct; .

Williams AH, Kim TH, Wang F, Vyas S, Ryu SI, Shenoy KV, Schnitzer M, Kolda TG, Ganguli S. Unsupervised Discovery of Demixed, Low-Dimensional Neural Dynamics across Multiple Timescales through Tensor Component Analysis. Neuron. 2018 Jun; 98(6):1099–1115.e8.

